# Unraveling a comparative landscape of protein-coding genes linked to neuroimmune function during adulthood consequent of prenatal alcohol exposure

**DOI:** 10.64898/2026.05.11.724451

**Authors:** Alissa Jones, Ariana Pritha, Ashlynn Aguilar, Andrea A. Pasmay, Justin Carter, Nikolaos Mellios, Shahani Noor

**Affiliations:** Department of Neurosciences, Reginald Heber Fitz Hall-145, MSC08 4740. University of New Mexico HSC, Albuquerque, NM 87131

**Keywords:** neuroimmune, prenatal alcohol, inflammation, proinflammatory, adult brain

## Abstract

**Background:** An overwhelming body of evidence suggests neuroimmune dysfunction as a key underlying mechanism of FASD-associated adverse CNS outcomes. While few studies have highlighted the lingering effects of prenatal alcohol exposure (PAE) on producing specific immune factors, others suggest a primed neuroimmune state in adulthood, in which a proinflammatory bias is unmasked following subsequent immune activation in later-life. However, the PAE-induced neuroimmune landscape in adulthood remains poorly defined. We hypothesized that PAE induces long-term changes in gene expression linked to neuroimmune function that may be brain region-specific.

**Methods:** Using long-read next-generation RNA sequencing of brain tissues from a previously established model of a moderate PAE in mice, we compared across six regions: medial prefrontal cortex (mPFC), anterior cingulate cortex (ACC), hypothalamus, hippocampus, midbrain, and medulla. A comprehensive bioinformatics analysis investigated PAE-induced changes, dysregulated gene pathways, and transcriptional regulators with a focus on neuroimmune function.

**Results:** Our data identified at least 60 differentially expressed genes per brain region, many of which were associated with neuroimmune function. Upregulation of multiple proinflammatory factors and pathways was observed, suggesting ongoing baseline neuroimmune activation, potentially involving PXR, TNF, TLR4, the complement pathway, and various cytokine and chemokine signaling. A comparative analysis identified multiple upstream transcriptional regulators across multiple brain regions, including MECP2, TCF7L2, and IL-4. Importantly, this unbiased analysis revealed heterogeneity across brain regions in the activation of canonical immune pathways and highlighted previously unprecedented roles of pathways such as PXR, matrix metalloproteases, and cytokine signaling (e.g., IL-15, IL-27, IL-17) in PAE.

**Conclusions:** PAE creates a unique inflammatory signature in the adult brain, even in the absence of secondary injury, with novel patterns of region-specific changes in genes implicated in glial-immune function. These data identified potential immune targets to elucidate the mechanisms underlying behavioral dysfunction and provide a framework for future therapeutic interventions.

## Introduction

Developmental exposure to alcohol has been increasingly recognized as a significant public health concern. Prenatal alcohol exposure (PAE) not only disrupts the developing brain but may also lead to structural and co-occurring functional changes that are long-lasting and extend beyond childhood and adolescence (Moore & Riley, 2015). Fetal Alcohol Spectrum Disorders (FASD)-associated behavioral and cognitive impairments lead to increased rates of depression, anxiety, mental health issues, learning disabilities, substance use, memory deficits, and sensory dysfunction in later-life (Moore & Riley, 2015). While current clinical data in FASD adults are sparse, a number of FASD-associated deficits persist into adulthood, increasing the health care costs over the life span (Tait et al., 2025). Understanding the molecular mechanisms underlying the long-lasting effects of PAE is important for developing effective healthcare strategies for individuals with FASD.

Findings from preclinical models have shed light on the long-term consequences of PAE, including impaired cognition and memory (Olguin et al., 2021), stress responses (Lam et al., 2018), and sensory dysfunction (Sanchez et al., 2017; Wang et al., 2022). These studies overwhelmingly suggest dysregulation of neuroimmune and neuroendocrine function as a major contributor to FASD-associated central nervous system (CNS) dysfunction (Smith, 2025). Cytokine production by CNS glia, such as microglia and astrocytes, not only regulates protective immune responses in the brain but also plays a critical role in brain development, maturation, and homeostatic neuronal function. The neuroimmune system regulates critical cellular mechanisms, including synaptic pruning and plasticity, cell survival, neurogenesis, mitochondrial function, and myelination (Imbriani et al., 2026). Enduring effects of PAE exhibit a proinflammatory immune bias in the CNS that extends beyond the embryonic period and early postnatal days (Baker et al., 2023; Ruffaner-Hanson et al., 2023; Ruggiero et al., 2018; Valenzuela et al., 2025). Few studies have suggested that PAE-induced changes are more prominent in adulthood than in adolescence (Baker et al., 2023). Thus, PAE-induced disruption of the neuroimmune system is suspected to be a central contributor to FASD-associated deficits.

PAE-induced proinflammatory immune bias during adulthood has been documented across various PAE paradigms varied in the duration, timing, and magnitude of alcohol exposure (Valenzuela et al., 2025), while heavy or binge drinking seems to be associated with profound neuroimmune changes (Cantacorps et al., 2017). However, these studies were largely conducted using targeted measures of classic proinflammatory cytokines such as interleukin (IL)-1β, tumor necrosis factor-α (TNF-α and IL-6, and discrete microglia and astrocyte activation markers. Even within this targeted analysis of discrete immune factors, variable responses were observed across brain regions (Gano et al., 2020), suggesting that PAE-induced long-term immune signature in the brain may differ not only across developmental ages, but also across brain regions. Extensive research indicates immune dysregulation in the hypothalamus and hippocampus, suggesting proinflammatory immune changes (Baker et al., 2023). Other reports included changes in the prefrontal cortex (PFC) and anterior cingulate cortex (ACC), regions that play a vital role in cognition, anxiety, and sensory dysfunction (Cantacorps et al., 2017; Hwang & Hashimoto-Torii, 2022). However, PAE-induced immune activation in other critical regions, such as the brainstem (e.g., midbrain and medulla), remains unknown. The midbrain includes important structures, such as the substantia nigra pars compacta and the ventral tegmental area, which play vital roles in regulating reward, motivation, working memory, and emotional processing (Bissonette & Roesch, 2016). PAE-associated behavioral deficits and dysregulation of dopaminergic activity involving these regions have been reported in clinical and preclinical studies in FASD (Shen et al., 1999).

A number of studies have now coined the term “immune priming” to describe a long-term consequence of PAE, characterized by heightened or maladaptive proinflammatory responses to *subsequent* CNS immune activation (Sanchez et al., 2017; Terasaki & Schwarz, 2017; Valenzuela et al., 2025) in adulthood. These studies proposed the idea that the PAE alters the “basal” state of the glial-neuronal gene expression, making it prone to proinflammatory immune function and later-life neurological dysfunction. PAE-induced immune priming has been demonstrated both with exogenous immune stimulation (e.g., lipopolysaccharide) and in sterile CNS injury models (e.g., physiological stress, nerve injury) (Baker et al., 2023; L. S. Terasaki & Schwarz, 2016). Increased levels of proinflammatory factors, including well-known cytokines such as interleukin-1β (IL-1β) and tumor necrosis factor-α (TNF-α), as well as various chemokines, have been reported in these studies. Several reports documented aberrant activation of the innate immune system in PAE via Toll-like receptor 4 (TLR4) and Nod-like Receptor, NLRP3 (Noor et al., 2023; Pascual et al., 2017; Ruffaner-Hanson et al., 2023). Similarly, anti-inflammatory cytokines appeared to vary, suggesting that the PAE-induced neuroimmune status reflects a complex interplay between pro- and anti-inflammatory molecules. Several studies reported subtle or no baseline changes of these select cytokines of interest, particularly in a low-to-moderate PAE paradigm, and only after the second hit or injury produced an altered neuroinflammatory outcome (L. S. Terasaki & Schwarz, 2016). Despite limited evidence of PAE-induced dysregulation of classic proinflammatory factors (e.g., IL-1β, TNF, and IL-6), a more comprehensive profiling revealed persistent PAE effects suggestive of an altered neuroimmune state at baseline, including dysregulation of anti-inflammatory molecules, other interleukins and microglial-activation mediators (Gano et al., 2020; Vella et al., 2025). While these targeted studies are highly informative, the magnitude and heterogeneity of baseline expression of the neuroimmune mediators that define PAE-induced neuroimmune reprogramming remain poorly understood. The current literature lacks a comprehensive, comparative investigation of the PAE-induced neuroimmune state in adulthood, which may explain the diverse neuroimmune outcomes of later-life challenges and vulnerability to CNS pathologies. To address this knowledge gap, we used next-generation long-read sequencing of tissue samples from a moderate PAE paradigm to analyze the mRNA transcriptome across six brain regions: cortical (mPFC, ACC), limbic (hypothalamus, hippocampus), and brainstem (midbrain, medulla). A comprehensive bioinformatics analysis was conducted to elucidate the most deregulated gene-expression pathways and molecular networks, and to identify transcriptional regulators that may underlie these PAE-induced changes. We compared various brain regions, with a focus on genes and pathways linked to neuroimmune function.

## Materials and Methods

### Animals

Animal procedures were permitted by the University of New Mexico Health Science Center’s Institutional Animal Care and Use Committee (IACUC) and in accordance with ARRIVE and National Institutes of Health’s guidelines for laboratory animal care and use (NIH Publications No. 8023, revised 1978). Animals were regularly monitored, including cage and bedding replacements every seven days.

### Prenatal alcohol exposure (PAE) paradigm

PAE and age-matched control mice were generated by the New Mexico Alcohol Research Center (NMARC) using a well-established moderate PAE paradigm (Brady et al., 2012, 2013; Olguin et al., 2021). Briefly, the Jackson Library provided C57BL/6J breeders; mice were habituated to a 12:12-hour reverse light/dark cycle and allowed free access to food (Teklad 2920X rodent chow) and water. Free access to either 0.066% (w/v) saccharin (SAC, non-PAE control mice) water or 10% ethanol sweetened with saccharin (PAE mice) was given for four hours daily (1000 to 1400 hr) during gestation. With blood ethanol concentrations of approximately 80 mg/dL, no significant changes in pup survival, weight, or litter size are observed with this paradigm (Brady et al., 2013). Offspring were weaned at 3 weeks and housed in groups of 2-4 mice per cage. PAE and control mice were housed in the same colony room and allowed to reach adulthood for experimental use. No more than one subject per litter was included to prevent potential litter effects. A total of twelve adult (7-8 months) female mice were used in this experiment (PAE n=6, control group n=6). Because of the lack of availability of age-matched adult male offspring at the time of the study, only female mice were included.

### Tissue Collection for RNA Extraction

Brain tissues from various regions were collected for RNA extraction. Mice were anesthetized with isoflurane (10 minutes at 5% vol./2.0% vol.). Mice underwent transcardial perfusion using ice-cold 0.1M phosphate-buffered saline (PBS; pH = 7.4; flow rate 10 mL/min). The brain was carefully removed from the skull. All brain regions were immediately dissected and flash-frozen in 1.5 mL DNase/RNase/Protease-free Eppendorf tubes (VWR International; Cat #: 20170-038) on dry ice. Tissues were preserved in a -80 °C freezer until RNA extraction.

### Total RNA Isolation

Tissues were homogenized using a motorized VWR disposable pellet mixer system. Per manufacturer’s instructions, RNA was extracted with Qiazol Lysis Reagent and miRNeasy Micro Kit (Qiagen; Cat #: 74004). The concentration and quality of total RNA were assessed using a NanoDrop spectrophotometer (Thermo Scientific, MA, USA).

### Profiling mRNA dysregulation in adult brain

RNA samples were sent to CD Genomics, where additional RNA quality assessments were performed, and RNA sequencing data were generated through next-generation sequencing technologies (NGS)-based long-read bulk RNA sequencing (Satam et al., 2023). Outliers were detected using Principal Component Analysis (PCA) and Euclidean distances between samples, and subsequently removed from the DE gene analysis, with no more than one outlier removed per brain region. Each dataset contained 5-6 biological replicates per group. Using R bioinformatics, data were then processed using the DESeq2 pipeline for all mRNA datasets, as described in our prior report (Pritha et al., 2026). DESeq2 is particularly useful for mRNA datasets, as it uses methods to reduce the number of false positives for genes that are lowly expressed or highly variable across samples. This yields an adjusted p-value (*p-adj*), which was used to assess the significance of the data. (Love et al., 2014) Datasets were processed using the adaptive shrinkage package (ASHR). This ASHR shrinkage method shrinks log fold changes (log FCs), thereby adopting a more conservative approach to statistical determination of effect size, reducing noise and potential false positives (Stephens, 2016). Results were filtered to only include protein-coding genes. Those with *p-adj* <0.05 and |log FC| > 0.25 were shown on the volcano plots.

### Analysis of gene expression pathways

All significantly differentially expressed genes (DEGs, *p-adj* < 0.05, no fold change threshold) from various regions were further analyzed with QIAGEN Ingenuity Pathway Analysis, or IPA (QIAGEN Inc., https://digitalinsights.qiagen.com/IPA) (Krämer et al., 2014). IPA is a machine learning algorithm that investigates interactions among DEGs to gain insights into biological pathways, functions, and critical molecular regulators that may play central roles in the underlying gene expression changes. Generation of all IPA networks and plots included all *p-adj-*significant DEGs. For any IPA analysis, a Z-score threshold of |2| is considered significant. Kyoto Encyclopedia of Genes and Genomes (KEGG) pathway analysis was performed using ShinyGo (KEGG version 0.85) (Ge et al., 2020). Gene ontology and KEGG enrichment analyses were performed separately for each brain region using the KEGG database. All mRNAs that had a log FC > |0.25| and a *p-adj* < 0.05 were included in the KEGG analysis.

## Results

### 3.1 PAE-induced DEGs in cortical regions

A distinct molecular profile with 46 upregulated and 152 downregulated significant DEGs (with a fold change threshold, log FC ≥ 0.25) emerged in the mPFC in adult PAE mice **(Fig. 1A).** PAE-induced upregulated DEGs involved diverse families of genes; including adhesion molecules (*Icam1, Cd72*), components of extracellular matrix (*Mmp11, Lama5*), immune regulators- *H2-DMa* (major histocompatibility complex II loading protein), *Ahnak2* (nucleoprotein), *Smad6* (a negative regulator of anti-inflammatory cytokine TGF-β), complement pathway related genes (*C1qa, Cqtnf4*) and genes related to cell death and oxidative stress response *(Chac1,* glutathione specific gamma-glutamylcyclotransferase 1) and *Nkx3-1*. Additionally, several genes linked to neuronal migration, synaptic function, excitotoxicity, and neuroendocrine signaling were dysregulated, including *Pcsk1n* (proprotein convertase subtilisin/kexin type 1 inhibitor), *Prr7*, and *Marveld1*. PAE downregulated a number of genes linked zinc finger family (*Zfp-965, -942, -943* and others), myelination (*Stag2*), RNA binding (*Qki, Syncrip, Hnrnph1, Hnrnpu*), protein degradation pathways (*Klhl15 and Klhl28, components of E3 Ubiquitin ligase*), mitochondrial function (*Cox20, mt-Nd3*) and cell survival and function (*Vbp1* and *Necab1*, calcium binding proteins). IPA analysis of the DEGs (*p-adj* <0.05) revealed dysregulation of zinc homeostasis pathways and multiple innate and adaptive immune pathways **(Fig. 4B).** Among the top 25 molecular networks, thirteen networks scored a Z-score of 22, while all 25 networks were scored a minimum score of 8 (Z-scores equal or greater than |2| are considered significant) (Supplemental File 1 contains all IPA networks for all brain regions). IPA molecular network analysis depicted activation of pro-inflammatory molecules like IL-1β (**Fig. 1C,** Z-score=22), complement signaling via C5AR1 and CD36 (Z-score=15). Although based on DEGs that displayed minimal fold changes (log FC <0.25), another top network (**Supplemental Fig. 1A,** Z**-**score =22) suggested activation of anti-inflammatory signaling (IL-10) in the mPFC, while TNF and TLR2 are predicted to be inhibited.

**Figure 1.**
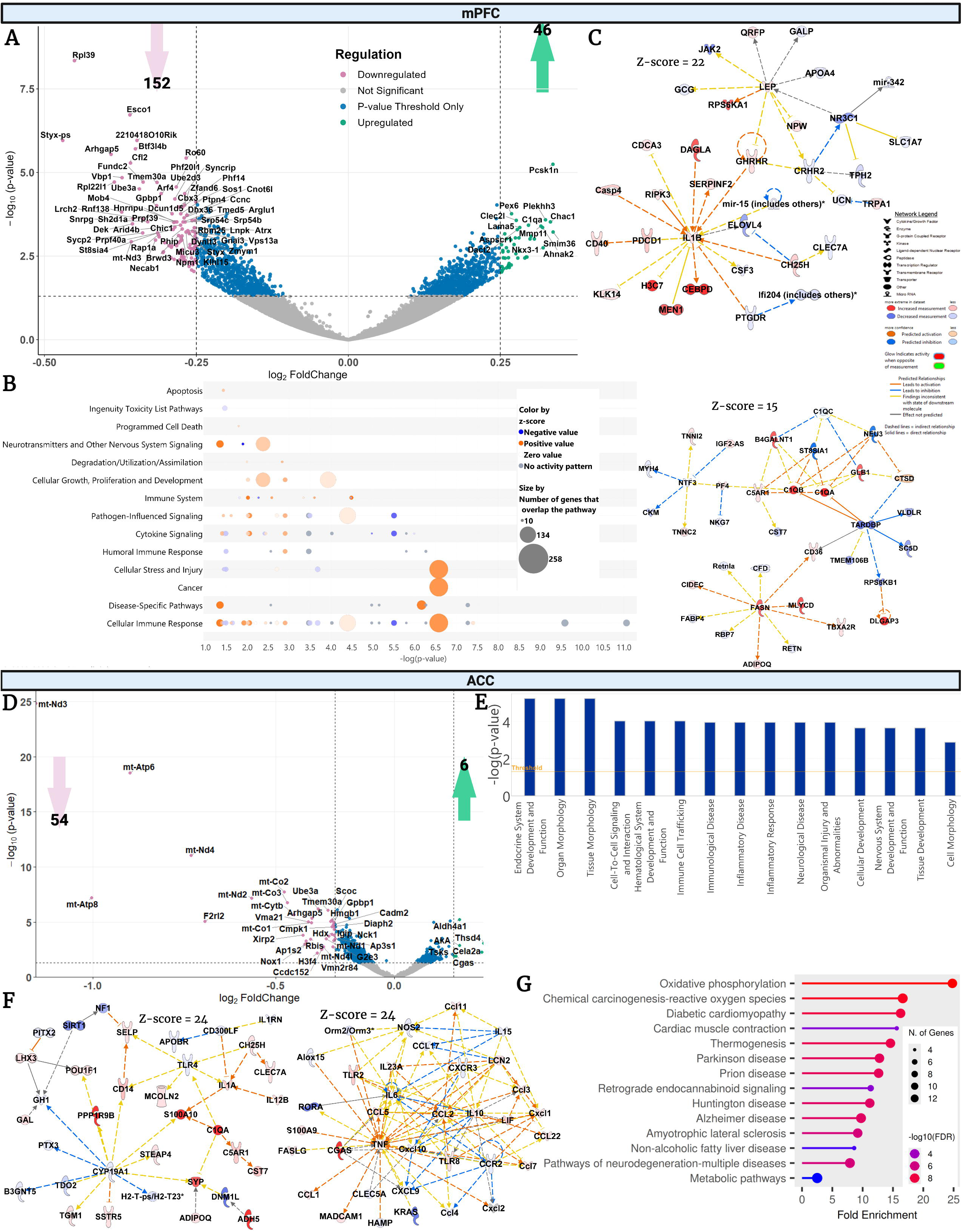
DE mRNAs and pathway analyses following PAE (A-D) mPFC: (A) The volcano plot displays significantly upregulated (green) and downregulated DEGs (pink) in response to PAE (|log FC| threshold = 0.25, *p-adj* < 0.05). (B) IPA canonical pathway bubble plot prediction based on *p-adj* significant mRNAs in the mPFC, with a brighter orange color indicating a higher Z-score for activation and a brighter blue color indicating a higher Z-score for inhibition of the pathway. Several cellular immune responses and cytokine pathways are predicted to be dysregulated. (C-D) IPA networks predict the activation of pro-inflammatory markers (IL-1β), histone subunits (H3C7), and transcription regulators (CEBPB), which are important for the epigenetic regulation of inflammation. Additionally, these molecular networks predict key components of the complement cascade (C1qa, C1qb) and microglial genes implicated in inflammatory activity (GLS1, CD14). **(E-I) ACC:** (E) DEGs (|log FC| > 0.25, *p-adj* < 0.05) displayed in the volcano plot as up- or downregulated. (F) Potential dysregulations in disease and biological functions based on *p-adj* significant genes in the ACC, with predicted changes in several immune-related pathways and diseases. (G-H) Molecular networks based on significant DEGs in ACC predict activation of pro-inflammatory cytokines and changes in cytokine and chemokine signaling. (I) Gene Ontology (GO) analysis predicts biological pathways dysregulated by DE genes linked to neurological diseases.

PAE elicited unique transcriptional alterations in the ACC, with 6 significantly upregulated genes and 54 significantly downregulated protein-coding genes **(Fig. 1D).** Strikingly, upregulated genes include *Cgas,* which emerged as a novel regulator of proinflammatory immune activation in the CNS. Other upregulated genes included *Cela2a* (elastase, linked to BDNF dysregulation) and *Aldh4a1*(aldehyde dehydrogenase). Among the PAE-induced downregulated DEGs, genes linked to mitochondrial function were predominant, including *mt-Nd3, mt-Atp8, mt-Atp6, mt-Nd4, and mt-Nd2, mt-Co2, and mt-Co3* all of which exhibited log FC ≤ -0.45. Interestingly, PAE-induced downregulation of *Nox1* (NADPH Oxidase 1) and *Hmgb1* was observed. IPA analysis of DEGs suggested the top fifteen diseases and biological functions, including “Endocrine system development and function”, “Immune cell trafficking”, “Inflammatory response”, and “Nervous system development and function” (**Fig. 1E)**. The IPA top network analysis revealed 25 molecular networks with a minimum Z-score of 9 **(Fig. 1F**. Among the top eight networks with a Z-score of 24, multiple networks displaying activation of cGAS signaling and its relations to a number of proinflammatory molecules, such as TNF, TLR-2, 8, and chemokines, Ccl1, Ccl2, Ccl7, and Cxcl1, were shown. Another top network showed upregulation of critical immune regulatory molecules, including C1q (complement signaling) and S100A10 (S100 Calcium Binding Protein A10), as well as IL-1 and IL-12 **(Fig. 1F)**. GO enrichment analysis suggested multiple pathways highly linked with mitochondrial dysfunction and respiratory complexes **(Fig.1G)**.

### 3.2. PAE-induced DEGs in limbic regions

Hypothalamic RNA expression exhibited a distinct gene expression pattern, with a large number of significant DEGs (2,510), with a threshold of *p-adj* <0.05 and a minimum of log2FC of 0.25. Of these, 1,471 genes were upregulated while 1,039 were downregulated **(Fig. 2A)**. Among the top upregulated 250 DEGs (all log2FC ≥0.43), a number of immune factors such as upregulation of *Mmp11* (matrix metalloproteinase 11), *Irf2bp1* (Interferon regulatory factor-2 binding protein), *Cox7c* (cytochrome oxidase C subunit) and *N4bp3* (NEDD4 binding protein 3), and *H1f2 and H1f2* (hypoxia inducible factors) were notable. Additionally, upregulation of adhesion molecules and cytokine and chemokine receptors was found (*Cd72*, *Cxcl17*, *Il11ra1*, *Il17rc)*. Upregulation of molecules involved in neuronal gene expression and plasticity (e.g., Neuronal PAS Domain Protein 1 (*Npas1*), *Nat8l*, solute carriers (*Slc4a2*), ATP-sensitive potassium channel (*Kcnj11*), *Pvalb* (parvalbumin, expressed in GABAergic interneurons), and *Vamp1* (component of SNARE complex in neurons) was found. PAE also induced genes related to neurovascular coupling (*Gja4*), astrocytic stress response (*Aldh3b1*), and oligodendrocyte differentiation (*Olig1*). PAE appears to disrupt the balance of multiple immune factors. Interestingly, downregulation of genes (log FC ≤ -0.7) linked to TNF signaling (*Traf1*, adaptor molecule of TNF receptor), mitochondrial function *(mt-Atp5*, *mt-Atp8*, *mt-Nd-3,-4,-2, 4l, mt-Cytb*), hemoglobin family genes (*Hba-a2,-a1*), and *Nlrp12* (NOD-like receptor) were evident. A reduction of Aquaporin (controls blood-brain barrier, *Aqp4)* gene expression was noted.

**Figure 2:**
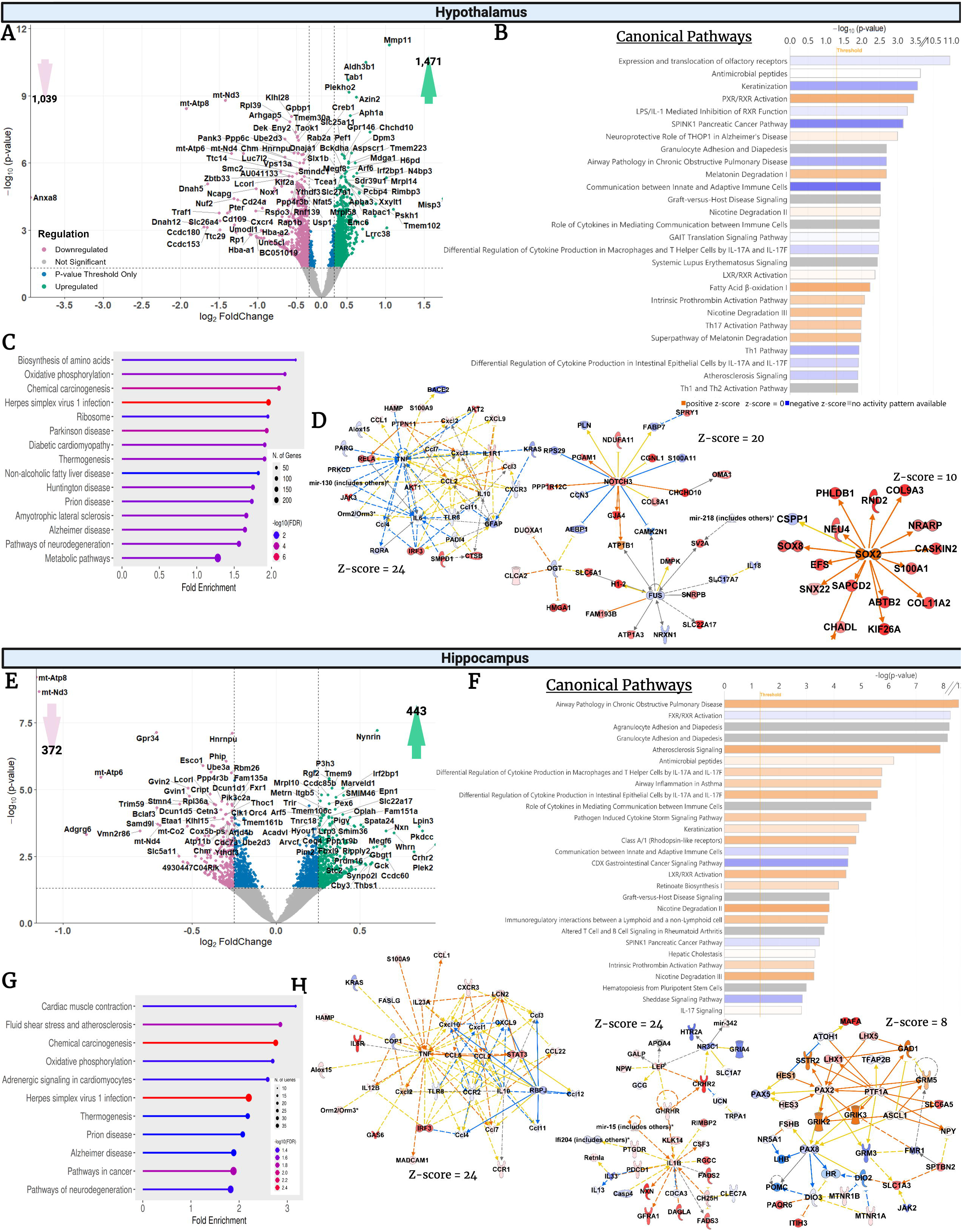
Limbic mRNA Dysregulation and Enriched Pathways due to PAE. (A-F) Hypothalamus: (A) Volcano plot displays significant DEGs with |log FC| threshold = 0.25 in PAE mice compared to controls. (B) IPA predicted canonical pathway dysregulation based on *p-adj* significant genes, with the same color scheme as described in Figure 1. (C) GO biological processes of DE-mRNAs predicted changes in oxidative phosphorylation and neurodegeneration. (D) Top molecular networks indicate predicted suppression of a number of pro-inflammatory factors like TNF and IL-18, strong activation of genes linked to myelination (SOX8, SOX2), and developmental disorders. **(F-J) Hippocampus:** (G) In the PAE hippocampus, 443 mRNAs were significantly upregulated (green) and 372 were downregulated (pink). (H) IPA predicted canonical pathway dysregulation based on *p-adj* significant genes, with the same color scheme as described in Figure 1. (I) GO biological pathway analysis of DE-mRNAs (J) Top molecular networks predict activation of proinflammatory cytokines, changes in endocrine system development and function, and molecular transport.

KEGG pathways and IPA pathways suggest dysregulation of oxidative phosphorylation, metabolic pathways, and immune pathways **(Fig 2B-C).** IPA networks predicted dysregulation of immunological pathways, particularly in the expression of chemokines, cytokines, and signaling molecules like CCL1, CCL2, CCL22, IL-10, IL12B, IL6, IL6R, TLR8, and TNF-α. A top IPA network predicted the suppression of TNF, TLR8, and GFAP, Il-18 **(Fig. 2D, Z-score =24, 20, 10)** while other networks predicted activation of SOX2, SOX8 and Notch 3, RelA (RELA proto-oncogene, NF-KB subunit), AKT1 (AKT Serine/Threonine Kinase 1) and IRF3 (interferon regulatory factor 3), critical factors involved in hypothalamic function and HPA and stress regulation, inflammation and myelination.

Substantial dysregulation of hippocampal mRNA expression was evident in PAE mice, with 443 upregulated and 372 downregulated significant DEGs **(Fig.2E)**. Among the upregulated genes, corticotropin-releasing hormone receptor 2(*Crhr2*) displayed the highest fold changes (log FC =0.79), a critical regulator of stress modulation, synaptic plasticity, and memory consolidation (Zheng et al., 2016). While a large number of genes displayed comparatively minimal fold dysregulations, many genes linked to proinflammatory immune function were evident in hippocampal DEGs. Upregulation of matrix metalloproteases (*Mmp -9, -14, -15*), chemokine receptor (*Cxcr4)*, cytokine receptor (*Il6ra)*, NFkBIE, *H2-Dma* (major histocompatibility complex), heat shock proteins (*Hspa1b*), *Atf6*, and *Irf2bp1* were found. Other molecular regulators of neuronal and glial survival and function, such as Wnt signaling (*Fzd7*, *Fzd8*) and growth arrest-specific proteins (*Gas1*, *Gas6*), are upregulated. PAE-induced downregulated DEGs with the highest fold changes included genes linked to mitochondrial function, such as *mt-Atp8*, *mt-Nd3*, *mt-Atp6*, *mt-Nd4*, and *mt-Co2*. Among other downregulated genes linked to immune function were *Gpr34* (G protein-coupled receptor 34), *Jak2* (Janus Kinase 2*), Il1rapl1* (IL-1 receptor accessory protein) and *Il-33*. Interestingly, molecules linked with a protective role in inflammation, such as *Nlrp6* (NOD-like receptor family pyrin domain containing 6,) *and Bcl2* (,B-cell lymphoma 2,)*, Htr2A* (serotonin receptor,)*, Nr3c1* (glucocorticoid receptor), were found to be downregulated in the PAE hippocampus. Other genes involved in glutamatergic neurotransmission, axon and dendrite growth, and synaptic stability (*Ncam2, Gria4, and Grm3*) were found to be downregulated in PAE hippocampus.

IPA predictions of canonical pathways included pathways such as pathogen-induced cytokine storm signaling, communication between innate and adaptive immune cells, and IL-17 signaling (**Fig. 2F).** GO biological processes predictions suggest DEGs impact pathways involved in oxidative phosphorylation, Alzheimer’s diseases and neurodegeneration pathways (**Fig. 2G**). IPA network analysis identified the top 25 molecular networks in the hippocampus; all networks had a Z-score of 8 or higher. One network (Z-score = 24) revealed a highly integrated molecular network with predicted activation of CCL1, CCL2, CCL5, IRF3, TNF, Alox15, MADCAM1, IL-6R, IFNG, IL-10, CCR1, IL-23a, but inhibition of CXCL10, CXCL9, TLR8, and CCL3 **(Fig. 2H)**. Two other significant IPA networks predicted activation of IL-1β pathway-linked immune molecules, as well as molecules linked to neuronal excitatory or inhibitory activity (e.g, GAD, Gria) (**Z-score = 24 and 8)**. These molecular networks show disrupted glial-neuronal communications critical for hippocampal function.

### 3.3. PAE-induced DEGs in brainstem regions

Our data from medulla tissue revealed immense differential regulation at a | log FC | threshold > 0.25 and *p-adj* < 0.05, with 1,109 significantly upregulated mRNAs and 1,309 downregulated mRNAs (**Fig. 3A**). With a more conservative threshold of | log FC | of 0.5, the medulla showed 196 total dysregulated genes (172 downregulated, 24 upregulated).The genes in the medulla spanned diverse biological functions, but a substantial proportion were associated with neuroimmune function. Among the DEGs, *Lyve1,* a marker of brain-resident macrophages, emerged as the top upregulated (log FC = 1.26). Other immune regulators, such *Cebpb* (transcription factor involved in proinflammatory immune activation), histone protein genes (*H3C-2*, *-3*, *-4*), *Nfatc4*, *Jund*, *Nfkbil1*, *Irf2bp1*, *Pcsk1n*, *C1qtnf4*, were notable with a log FC ≥ 0.5 (J. Li et al., 2025). Other upregulated DEGs with links to immune function included cytokines and their receptors, *Il21r*, *Il11ra2*, and *Il34*. Downregulated DEGs with at least log FC ≥ |0.5| change included genes linked to mitochondrial gene function (*mt-Atp6*, *-8*, *mt-Co3*, *mt-Nd2*) and various DEGs linked to zinc finger family protein, *Nox1* and *Gpr34*, *Cd36* and *Cox20*. IPA analysis predicted several different immune pathways (**Fig. 3B**), including activation of TNF and IL-6 (as well as MyD88, Cebpb, Jak3, Akt1-2, RelA, and Tmem119), indicative of a pro-inflammatory state in the medulla (**Fig. 3C,** Z-score =15). Another network highlighted the involvement of aberrant neuronal activity, potentially via activation of GRIN1 and GRIK5 (**Supplemental Fig. 1B**, Z-score =15).

**Figure 3:**
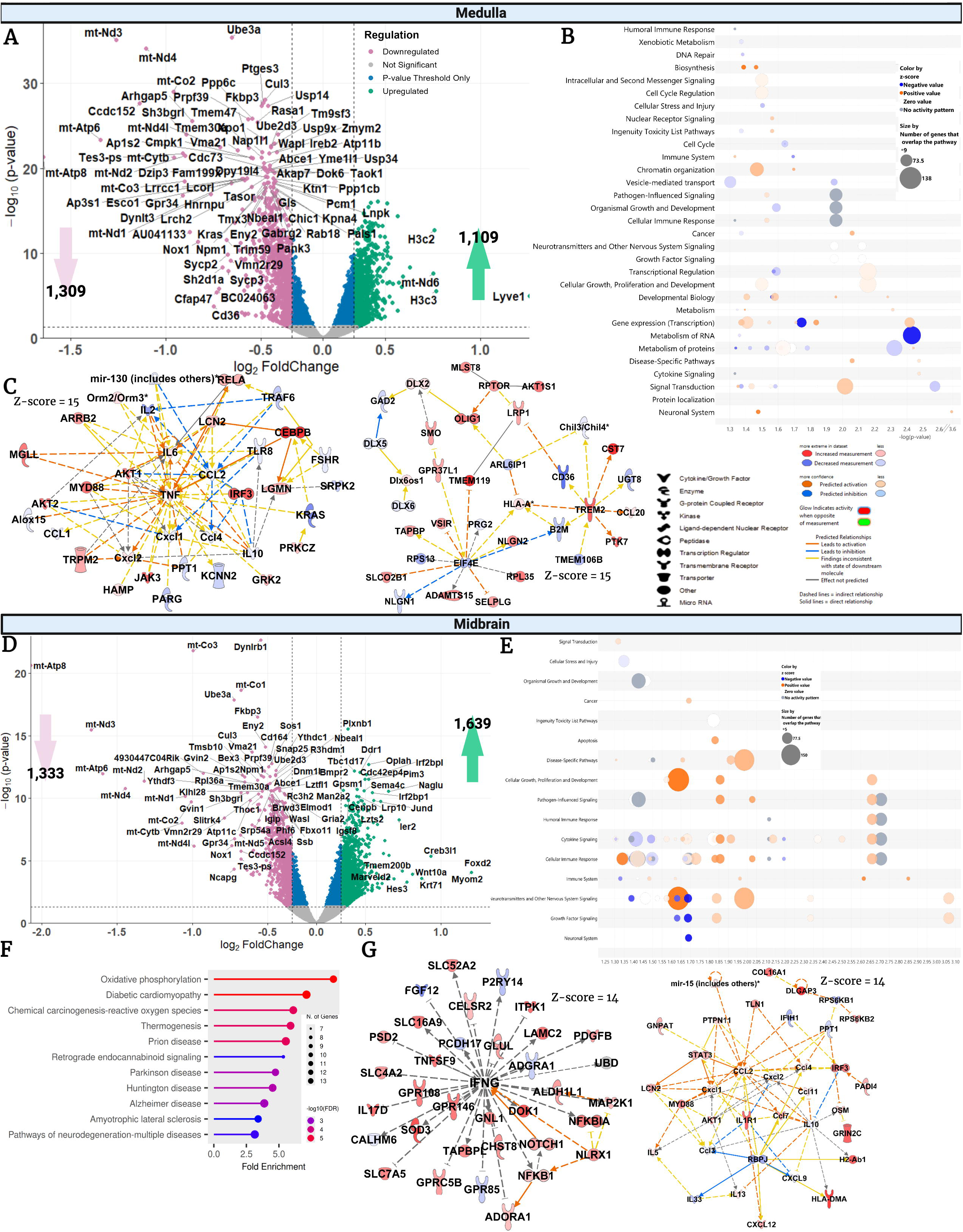
Brainstem mRNA Dysregulation and Pathways. (A-F) Medulla: (A) Volcano plot depicting DE-mRNAs in the PAE medulla. A total of 1,109 mRNAs were significantly upregulated and 1,309 were downregulated (*p-adj* < 0.05, |log FC| > 0.25). (B) IPA canonical pathway bubble plot prediction based on *p-adj* significant mRNAs in the medulla, with a brighter orange color indicating a higher Z-score for activation and a brighter blue color indicating a higher z-score for inhibition of the pathway. Bubble size increases with the number of genes contributing to the pathway. (C) IPA network 1 of the medulla (z-score = 15) predicted upstream regulators and key molecular networks altered in the medulla, including activation of TNF and IL-6. (D) IPA network 12 of the medulla (z-score = 15) predicts changes in cell morphology, development, growth, and proliferation. **(E-I) Midbrain mRNA.** (E) Volcano plot of DE-mRNAs in the midbrain of PAE versus SAC mice. A total of 1,639 mRNAs were significantly upregulated (green) and 1,333 downregulated (pink) (*p-adj* < 0.05, |log FC| > 0.25). (F) IPA canonical pathway bubble plot prediction based on *p-adj* significant mRNAs in the midbrain (same parameters as panel B). (G) GO biological processes predict pathways involved in oxidative phosphorylation and neurological disorders. (H) Top networks revealed predicted dysregulation of molecules in inflammatory response (NFKB1, IL17D) and activation of several pro-inflammatory molecules.

The midbrain showed even greater PAE-induced transcriptional perturbation, with 1,639 upregulated and 1,333 downregulated mRNAs in PAE **(Fig.3D).** With a more conservative threshold of | log FC| of 0.5, the midbrain contained 188 dysregulated genes (67 upregulated, 121 downregulated). Of the genes linked to neuroinflammation, upregulation of interferon-related genes (*Ifi2712b*, *Irf2bp1*, *Irf3*), *Jund*, *Gas1*, *Il17rc*, *Tnfrsf4,* and adhesion molecules (*Itga11*) showed a minimum log FC of 0.5. Upregulation of other integrin molecules (*Itgal, Itga7*) and proinflammatory *Il16* and *Il1r1* was significantly increased in PAE. Interestingly, various CCL21 isoforms (*Ccl21a*, *Ccl21b*, and *Ccl21d*) and *Ccl19* were all upregulated in PAE midbrain. These chemokines regulate neuron-glia communication, neurodegeneration, and behavioral deficits (Leser et al., 2025). *Wnt10a* was also upregulated, a gene linked to astrocyte activation, neuroprotection under oxidative stress (K. Li et al., 2024). Anti-inflammatory cytokines (*Il2*, *Il4*), along with genes linked to metabolism and energy and mitochondrial function (*mt-Atp-6, -8*, *mt-Nd-2,-4,-1*, *mt-Cytb*, and *mt-Co1,2,*3 and *Nox1*), were found to be downregulated due to PAE. Dysregulation of molecular regulators associated with neuronal activity during inflammation, including *Ier2* (Immediate early response 2) and *Il1rapl1*, was observed (Montani et al., 2017).

IPA analysis suggested that these DEGs were significantly enriched for genes regulating cytokine signaling, neurotransmitter signaling, and cellular immune responses (Fig.3E), with predictions of both “activation” and “inhibition” of immune-cell signaling cytokines and chemokines. GO enrichment analysis predicted changes in pathways such as oxidative phosphorylation and retrograde endocannabinoid signaling (Fig. 2F). All top 25 networks exhibited a strong Z-score of 14, with 35 focus molecules. These networks predicted activation of a number of molecular regulators of proinflammatory function, including IL-1R, IRF3, CCL2, NFKB, Myd88, NLRX1 (Nod-like Receptor), Icam-1, as well as the complement pathway (**Fig. 3G; Supplemental Fig. 1E-F**).

### 3.4. Overlapping DEGs linked to immune functions across multiple brain regions

Further analysis of significant DEGs (*p-adj*<0.05, log FC ≥ 0.25) revealed a number of both unique and overlapping genes in two or more regions in the PAE brain. **Figure 4A** shows the number of DEGs common in various regions; a detailed list of these DEGs is found in **Supplemental File 2**. In **Table 1**, selected overlapping genes known to play immune regulatory functions are listed with their fold changes. Of interest, several mitochondrial genes were dysregulated across all analyzed regions, except the mPFC. These include *mt-Co1*, *mt-Apt8, mt-Atp6, and mt-Nd4*, all of which were downregulated in the majority of the regions analyzed.

**Figure 4.**
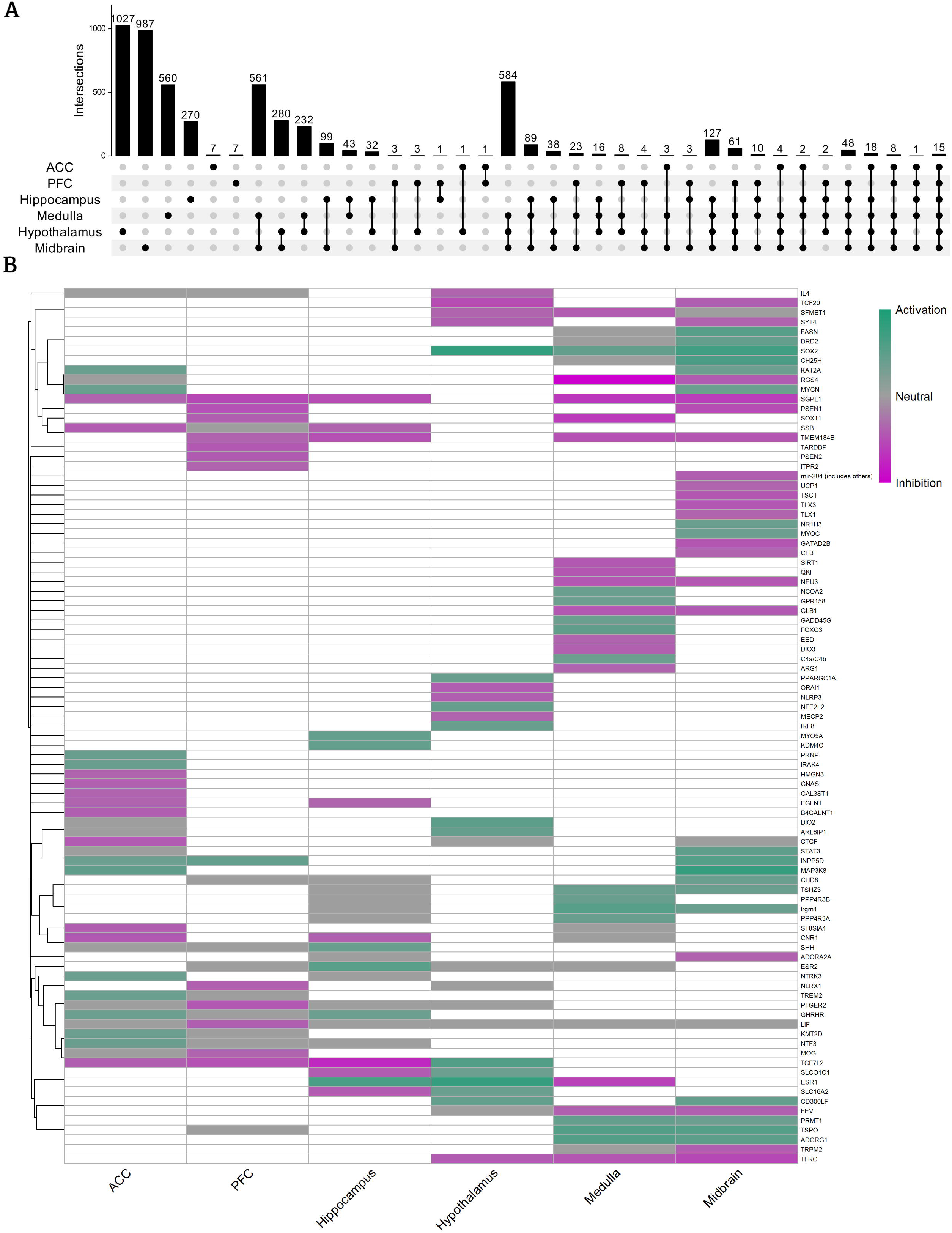
Convergent and Region-Specific Transcriptional and Regulatory Alterations Across the Brain. (A) UpSet plots showing the number and overlap of DE mRNAs (|log FC| < |0.25|, *p-adj* < 0.05) across the six brain regions analyzed. Vertical bars represent the number of overlapping genes in each region or region combination, while horizontal bars display the total number of DE genes in each region. (B) Heatmap of IPA-predicted upstream regulators across brain regions. Darker purple indicates stronger predicted activation, while darker green indicates stronger predicted inhibition (IPA Z-score).

**Figure 5.**
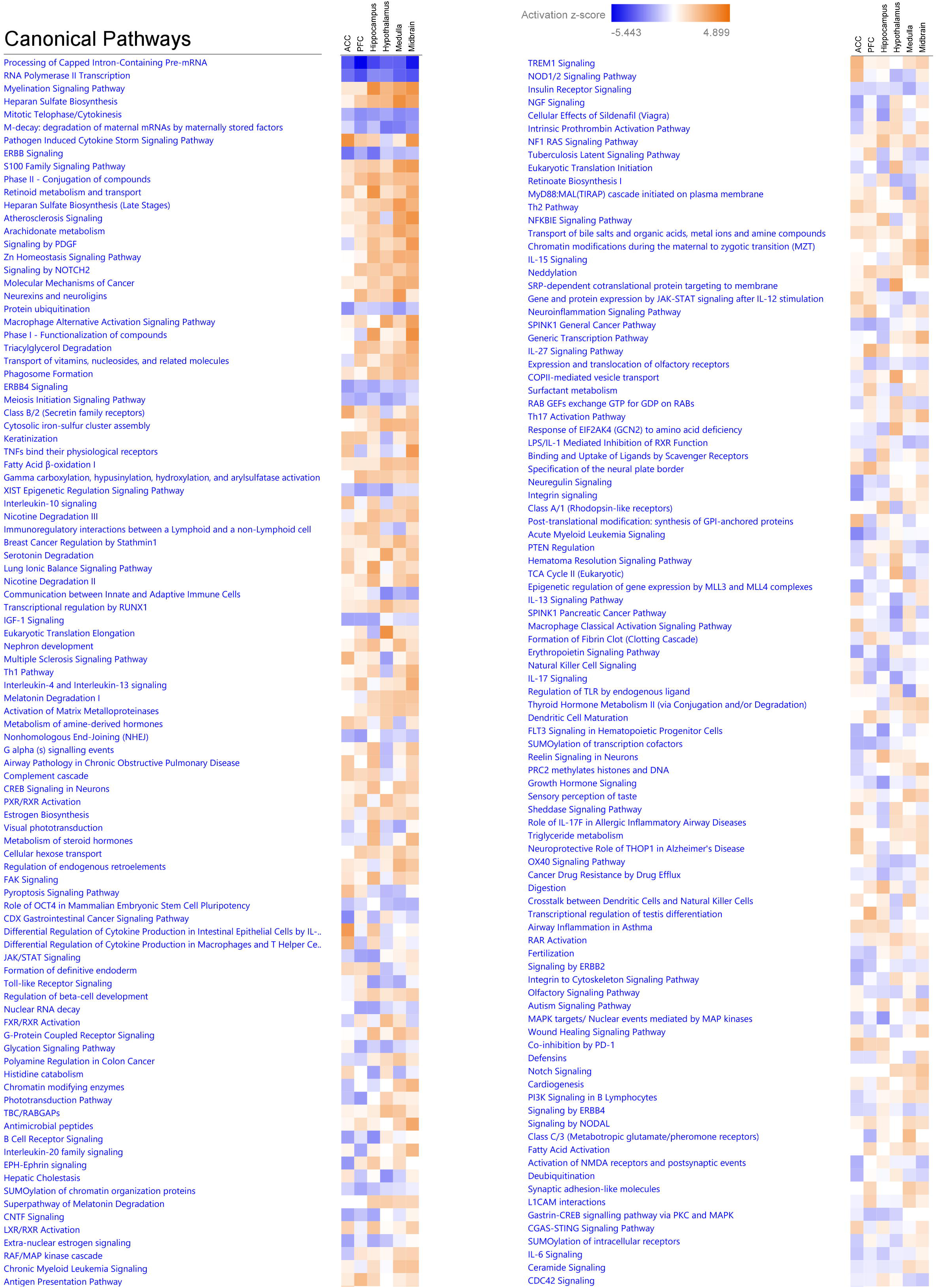
Canonical Pathway Heatmap Across Brain Regions. Heatmap of the number of overlapping canonical pathways (from IPA analysis) across brain regions, based on *p-adj* significant mRNAs. Pathway overlap is visualized using color intensity, with positive (orange) and negative (blue) Z-scores.

**Table 1.**
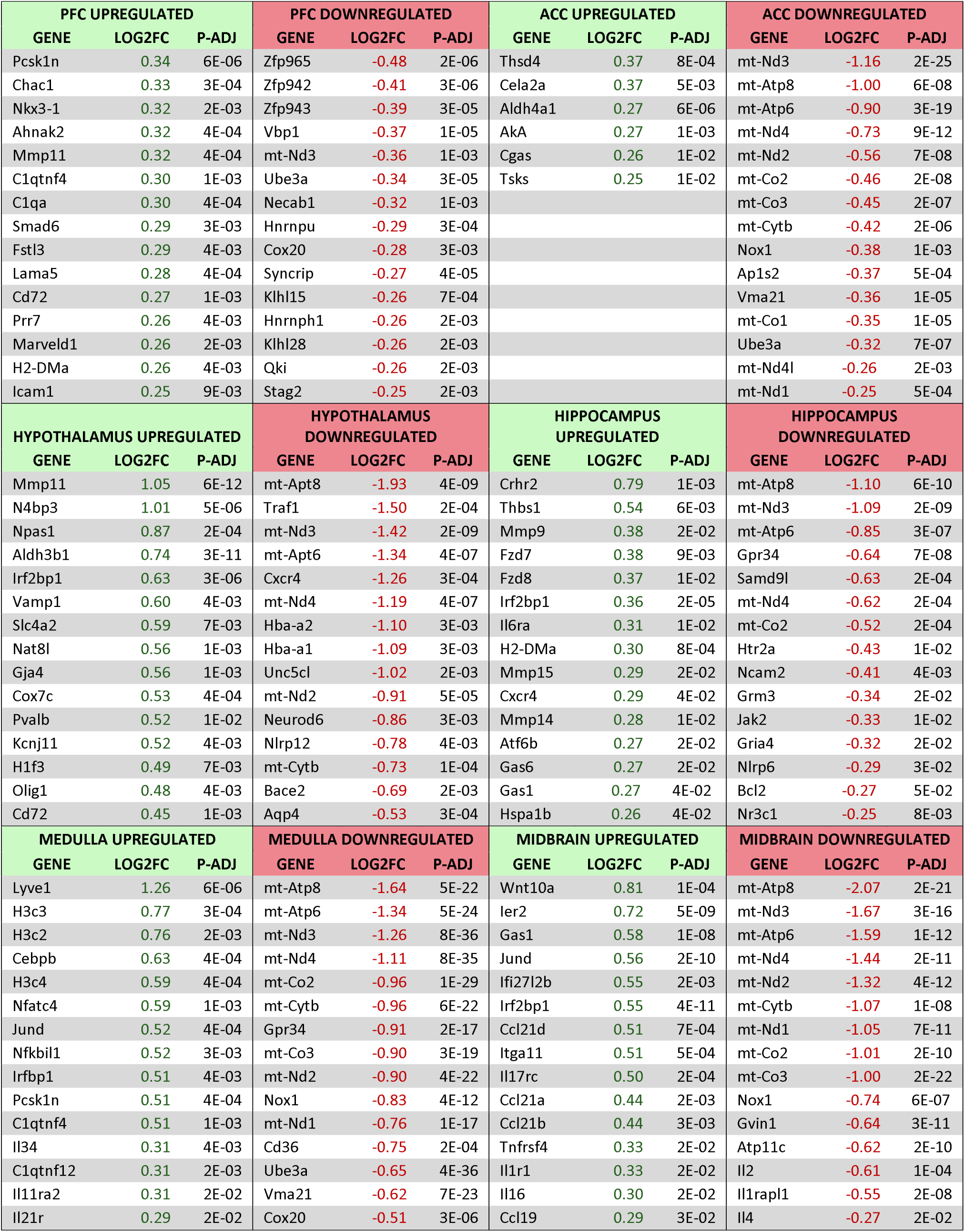
Top significant DEGs (with a minimum of log2FC of 0.25, selected from the top 15) linked to neuroimmune function

**Table 2:**
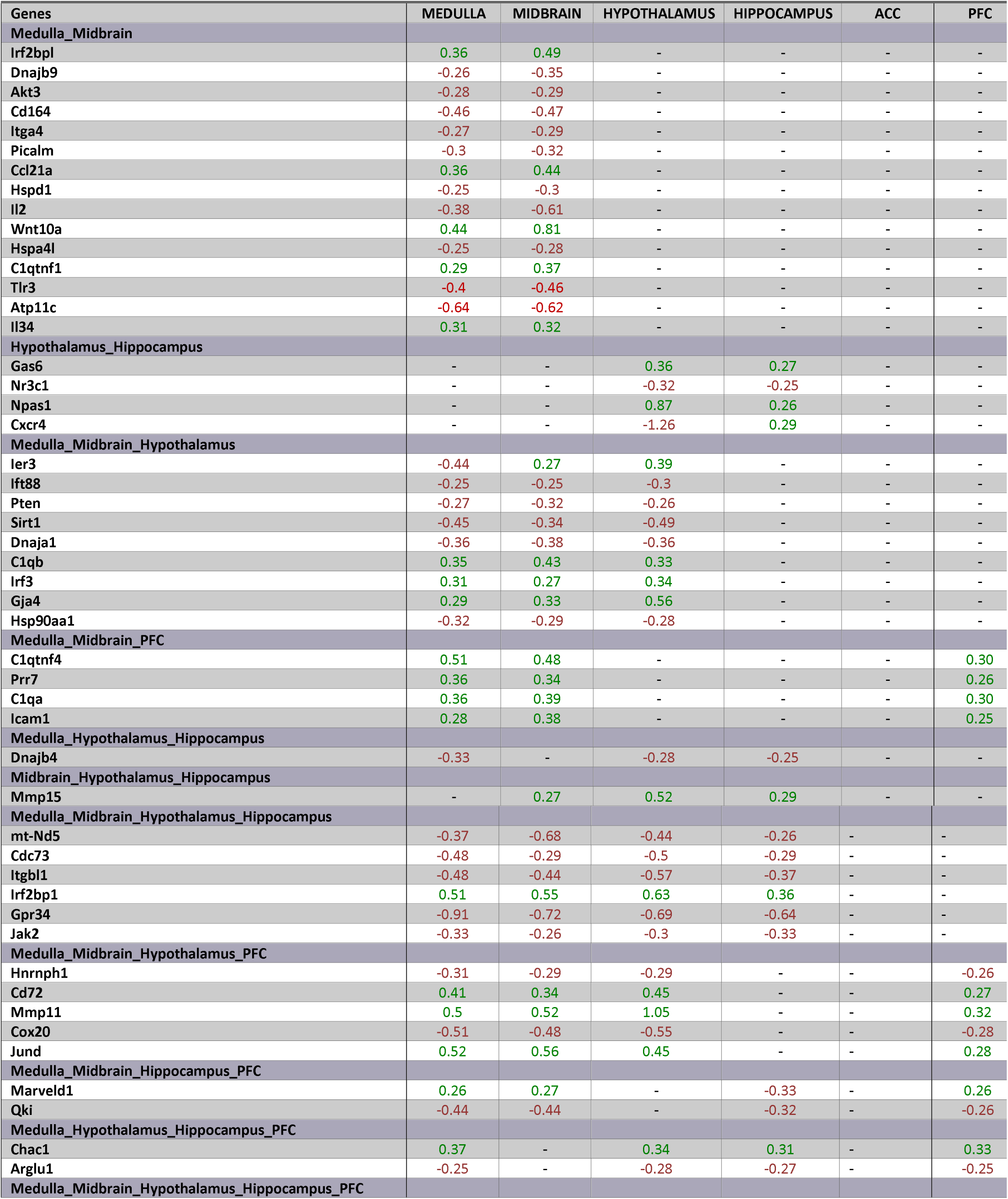
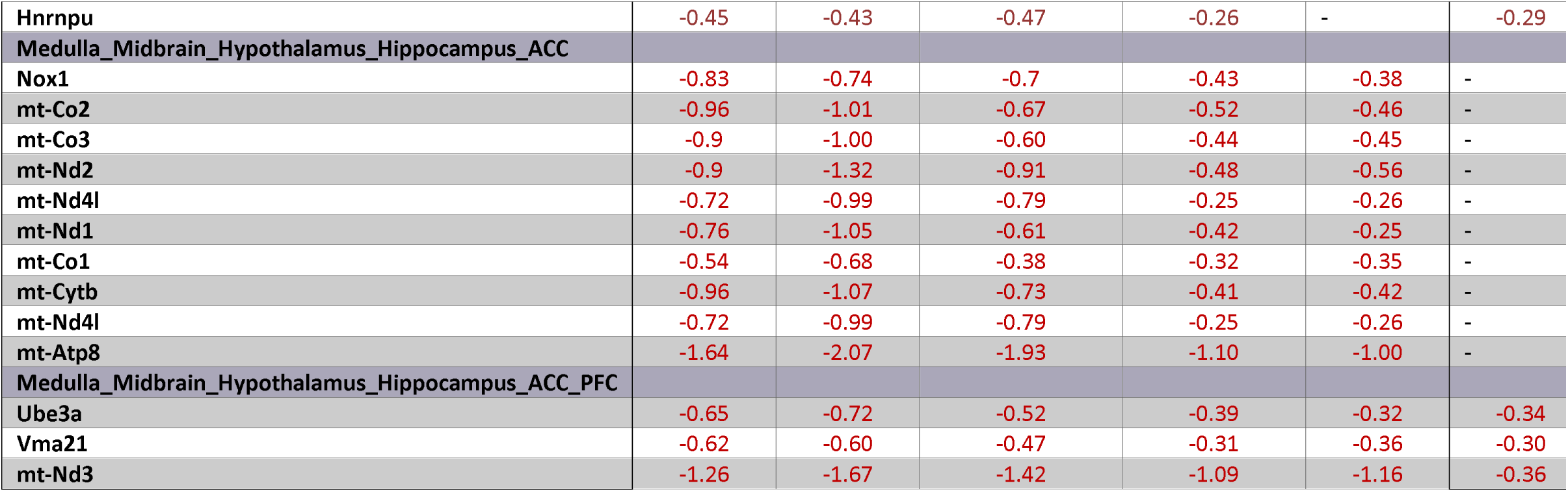
Selected overlapping significant DEGs across brain regions. Log_2_FC s are shown each significant DEGs in the indicated brain region. Blank cells indicate that the gene did not meet dysregulation criteria in that region (log_2_FC > |0.25| *p-adj* < 0.05).

In the brainstem regions and the mPFC, complement proteins *C1qa* and *C1qtnf4*, and the adhesion molecule *Icam1*, were upregulated in PAE. Matrix metalloprotease proteins (*Mmp11*, *Mmp15*) were found to be upregulated in at least three brain regions analyzed. Notably, the proapoptotic factor, *Chac1,* was upregulated in four brain regions. Although the roles of *Sirt1* and *Pten* are context-specific, their anti-inflammatory effects are well established, and these molecules are downregulated in brainstem regions and the hypothalamus. Downregulation of IL1RAPL1 was notable in the brainstem, as well as limbic regions. PAE-induced changes in mRNA transcripts shared a considerable overlap in brainstem regions; 561 mRNAs were found in common in the midbrain and medulla. Common to both regions, several proinflammatory molecules, such as *Ccl21*, *Wnt10a*, *C1qtnf1*, *Nfatc4* (Nuclear factor of activated T-cells 4), and *Shank3* (SH3 and multiple ankyrin repeat domains 3), were found upregulated. In contrast, proteins with anti-inflammatory or neuroprotective roles, such as *Il2*, *Picalm* (phosphatidylinositol binding clathrin assembly protein), *Hpgd* (Sun et al., 2025)), *Atp11c*, and *Ak3*, were downregulated. However, other genes shown to be involved in innate immune activation, such as *Hspa4l*, *Dnabj9*, *Tlr3*, and *Hspd1*, were also downregulated in brainstem regions. Interestingly, fifteen mRNAs were consistently identified as downregulated across all six regions, including *mt-Nd3, Ube3a* (Ubiquitin ligase), *Vma21,* and zinc finger proteins *(Zfp931, Zfp943, Zfp948, and Zfp938).* Similarly, downregulation of RNA-binding and transcriptional and post-transcriptional regulatory molecules, such as *Hnrnpu, Qki,* and *Hnrnph1,* was observed in the majority of these regions **(Table 1).**

### A comparative analysis of upstream regulators in various brain regions

IPA analysis of upstream regulators was conducted to identify potential master regulators of gene-expression changes in these datasets. The heatmap displays the presence of any upstream regulator with a significant Z-score (defined as ≥ |2|) in at least one dataset, along with their predicted activation or inhibition based on the significant (*p-adj*<0.05) DEGs in the corresponding dataset (**Fig. 4B**). This analysis revealed multiple immune regulators with significant Z-scores across two or more regions, whereas in certain brain regions, unique upstream regulators were also observed. IL-4 was significantly inhibited in the hypothalamus, whereas TREM2 and IRAK4 activation were observed in the ACC. Notably, potent immune modulators, such as INPP5D and MAP3K8, were predicted to be activated in the cortex and brainstem. TCF7L2 displayed contrasting effects in cortical regions and in the hippocampus and hypothalamus. Estrogen receptor (ESR1) was predicted to be activated in the hippocampus and hypothalamus. These data identified the diversity of potential immune regulators potentially implicated in the PAE brain.

### A comparative analysis of dysregulated canonical pathways in various regions

A comparative analysis of canonical pathways associated with DEGs across brain regions revealed striking similarities in the activation or inhibition of multiple pathways implicated in immune and critical neuronal functions. Top 190 pathways (ranked by z-score) included TNF binding to physiological receptors, pathogen induced cytokine storm signaling, IL-10 signaling, IL-4 and IL-13 signaling, Th1, activation of matrix metalloproteinases, communication between innate and adaptive immune pathways, complement signaling and PXR/RXR activation, TLR and TREM1 signaling and NOD1/2 signaling pathway, cytokinesis, IL-15 and IL-27 signaling, Jak/stat signaling and integrin and pyroptosis signaling pathway. Although some brain-region-specific activation patterns were evident in these pathways, complement cascade activation and other proinflammatory signaling pathways (TREM1, TNF, Th1, IL-15) showed profound activation in the cortical regions, and hippocampus. A comparatively weaker IL-10 signaling was evident in the hypothalamus, whereas IL-4 and IL-13 were predicted to be stronger in the mPFC and midbrain. Other pathways linked to general RNA and protein turnover are also noted, including the processing of capped intron-containing pre-mRNA, RNA polymerase II transcription, and protein ubiquitination. Interestingly, pathways linked to neuronal processes and oligodendrocytes were also depicted, including myelination, Zn homeostasis, CREB signaling in neurons, NOTCH signaling, serotonin and nicotine degradation, MAP kinase signaling, and growth factor (IGF and NGF) signaling and nephron development.

## Discussion

Numerous studies demonstrate that early life adversity predisposes an individual to a primed glial-immune state during adulthood that dictates vulnerabilities to develop neurological dysfunction, including psychiatric disorders, neurodegeneration, autoimmune disorders, and influences healthy aging. While the long-term effects of PAE are increasingly being recognized as a risk factor for developing chronic CNS dysfunction in adulthood (Vella et al., 2025), current literature lacks a better understanding of the PAE-induced neuroimmune landscape. While prior studies suggest that the effects of PAE on neuroinflammatory cytokine production may diminish over time for select immune mediators, other studies have provided clear evidence of neuroimmune activity (Vella et al., 2025). To our knowledge, this is the first unbiased characterization and comparative analysis across brain regions of the enduring effects of PAE in this age range. PAE induces distinct neuroimmune changes across discrete brain regions. While an inflammatory tone is evident across most regions and overlapping immune factors were identified, PAE-induced changes appear to involve a complex dysregulation of both anti- and pro-inflammatory immune mediators, potentially involving diverse transcriptional regulators and immune pathways.

Prior research investigating transcriptomic dysregulation in PAE reported both short-term and long-term changes in various brain regions, including the postnatal hippocampus (Lunde Young et al., 2019), postnatal cerebellum (Holloway et al., 2023), embryonic neural tube (Boschen et al., 2020), mouse cortex (Mishra et al., 2023; Sambo et al., 2022) and olfactory bulb (Gano et al., 2020). Our data are consistent with substantial evidence of immune activation in cortical areas and the hippocampus in the adult brain due to PAE (Baker et al., 2023; Gano et al., 2020; L. S. Terasaki & Schwarz, 2016). Our data suggest upregulation of *Pcsk1n*, complement activation, and molecules linked to Wnt signaling, as well as IL-10 signaling, in multiple brain regions. Although observed in studies on early postnatal days, PAE-induced dysregulation of these factors has also been reported in other preclinical studies (Holloway et al., 2023; Lunde Young et al., 2019; Sambo et al., 2022). Importantly, a basal level of proinflammatory immune activity was revealed in brainstem regions. Our data are suggestive of involvement of chemokine (*Ccl21*), Wnt (*Wnt10*), and complement (*C1qtnf1*) pathways, and of downregulation of the immunosuppressive phenotype (CD164, IL-2), along with concurrent changes in the neuron-enriched potassium channel *Kctd9*, an emerging regulator of various neurological diseases and neurodegeneration (Teng et al., 2019). IL-6 was upregulated in the brainstem, consistent with findings in other brain regions in PAE adulthood (Baker et al., 2023). Together, these data highlight a previously unexplored effect of PAE on neuroimmune activation in the brainstem, which may contribute to PAE-associated dysfunction.

Mitochondrial cellular respiration plays a critical role in regulating cellular functions in both neurons and glia. Prior studies have reported as early as the immediate effects of fetal alcohol exposure, including downregulation of mitochondrial complexes I and V, linked to cognitive impairment (Terracina et al., 2024). Strikingly, we observed consistent downregulation of mitochondrial genes across various brain regions. These data indicate that perturbations in energy balance and oxidative stress appear to be primary mechanisms of the lingering effects of PAE, which links to neuroinflammation. Mitochondrial damage, often as a result of oxidative stress, can contribute to neuroinflammation via compromising the blood-brain barrier (BBB) (Sadikot et al., 2019). Several molecules, such as *Aqp4*, *Gpr34*, and glucocorticoid receptors, were found to be downregulated by PAE, which may impair neuroinflammatory signaling, synaptic plasticity and microglia-mediated myelin debris sensing (Lin et al., 2023). Several brain regions displayed upregulation of matrix metalloproteinases, which are also essential for synaptic plasticity and BBB integrity, including modulation of ROS species and proinflammatory immune activation (Bitanihirwe & Woo, 2020). Interestingly, *Nox1* was found to be downregulated in most brain regions, a key enzyme that generates superoxide, a reactive oxygen species. Notably, in cortical regions, *Hmgb1* was downregulated. *Hmgb1* has been implicated in chronic alcohol exposure in adulthood (Crews et al., 2013) and other preclinical studies with upregulation of *Hmgb1* mRNAs in early postnatal days in PAE females (Ruffaner-Hanson et al., 2023). While downregulation of *Hmgb1* and other factors, such as TNF, may indicate reduced potential to activate proinflammatory immune signaling via NFKB-mediated transcription, other immune activators, such as *Cgas*, have been observed to be upregulated in cortical regions, which may increase the cGAS-STING inflammatory pathway innate immune axis (Barrett et al., 2021), as predicted by our canonical pathway analysis. *Nr3c1* (Glucocorticoid receptor), a critical receptor that balances pro- and anti-inflammatory pathways (Kim et al., 2020), was found to be downregulated in the PAE hippocampus.

Lastly, although our study focused on neuroimmune genes, a number of molecular regulators of neuronal function (e.g., neuronal channels, solute transporters) were identified, suggesting altered glial-neuronal interactions consequent to PAE, which set the stage for neurobehavioral and neuroimmune outcomes following later-life exposures. Moreover, dysregulations of components of mitochondrial complex 1, protein degradation pathways, zinc finger proteins, and interferon regulatory factors and IL1RAPL1 have been linked to other neurodevelopmental disorders and mental disabilities (Furumai et al., 2019; Montani et al., 2017). Interestingly, although the role of *Vma21* in FASD remains unknown, *Vma21* promotes the reversal of cognitive deficits (Wu et al., 2024), a neuromodulator found to be downregulated across all PAE brain regions. Similarly, IL1RAPL1 mediates excitatory synapse formation, influencing PSD-95 and other synaptic scaffolding proteins (Liu et al., 2019).

Multiple transcriptional regulators appeared to be involved in PAE-induced transcriptomic changes, with overlapping candidates identified; many were unique to specific brain regions. Although the exact role of TCF7L2 remains unknown, it is among the most dysregulated across regions, with contrasting effects on predicted activation or inhibition in specific regions. Interestingly, TCF7L2 has been shown to activate the TLR4 and beta-catenin pathways, which have been extensively implicated in multiple reports of PAE (Noor et al., 2023; Ruffaner-Hanson et al., 2023). Similarly, several other critical transcriptional regulators of neuroinflammations, including MECP2, were dysregulated in the hypothalamus. SOX2 is crucial in the maintenance of the HPA axis (Jayakody et al., 2012). Our data were suggestive of SOX2 activation in the hypothalamus and other brainstem regions. Together, these data provide evidence of basal dysregulation of critical molecular regulators that may predispose primed immune responses to PAE conditions.

Our findings elucidate potential dysregulation of diverse canonical pathways implicated in immune dysfunction in the adult brain consequent of PAE. Several immune pathways linked to microglia proinflammatory function were identified, including complement cascades, Wnt signaling, and PXR and TREM1. Interestingly, our data projected activation of PXR/RXR in the majority of regions of the PAE brain. Notably, recent studies indicate that PXR regulates innate immune responses by promoting NLRP3-mediated inflammatory signaling (Hudson et al., 2019). Although our data do not provide evidence of activation of TLR4-derived proinflammatory cytokines consistently in all regions at baseline (in the absence of secondary immune stimuli), our findings indicate potential alterations in multiple proinflammatory chemokine systems and in cytokine activity (TNF, IL-17, IL-20, IL-15) across these regions. Consistent with the literature, an increase in IL-10 was suspected from our DEGs and pathway analysis, potentially to mitigate ongoing basal immune proinflammatory function. Transcriptional changes in other pathways (e.g., IL-4 and fatty acid oxidation) linked to anti-inflammatory and immune-regulatory signaling were observed. A number of these innate and adaptive immune pathways are highly interconnected. While few studies have used targeted approaches to confirm the key contributions of certain pathways (e.g., TLR4, NLRP3) (Noor et al., 2023; Pascual et al., 2017; Sanchez et al., 2017), the roles of other pathways in driving neuroimmune dysfunction in adulthood remain to be explored.

### Limitations

Sex is a key variable in preclinical and clinical research on FASD (Mishra et al., 2023). PAE may generate sex-specific changes in gene expression in brain regions. While this study focused on female mice, our ongoing and future research will validate sex-specific immune factors of interest in both sexes. Although several observations from these datasets were consistent with protein-level changes reported in prior studies (Sanchez et al., 2017; L. Terasaki & Schwarz, 2017; Vella et al., 2025), given the nature of bulk RNA sequencing data, the contributions of various glial cell types to immune changes require further validation at the protein level.

### Conclusion

PAE leads to numerous gene-expression changes associated with neuroimmune function in adulthood. PAE-related long-term glial immune changes are diverse and show a distinct pattern in specific brain regions. PAE-induced immune reprogramming in these brain regions appears to be an intricate balance of pro- and anti-inflammatory factors, concurrent with changes in molecular regulators of neuronal function. Together, this study provides a comprehensive overview of the heterogeneity of PAE-induced effects, identifies novel immune molecules and pathways, and transcriptional regulators, and elucidates their unprecedented involvement in PAE-associated dysfunction. These data provide a critical framework for future explorations of targeted immunomodulatory therapies in FASD.

## Acknowledgements

We thank Dr. C. Fernando Valenzuela, Minerva Murphy, and Diane Jimenez from the New Mexico Alcohol Research Center (NMARC) for providing PAE and control mice. Formatted figures were created with BioRender. Jones, A. (2026) https://BioRender.com/hl0adxj

## Author Contributions

SN and NM designed and conceptualized the study. ANJ, ANP, and SN wrote the initial drafts. NM, AAP, and ANP edited and contributed intellectually to the interpretation of the data. AAP and JRC conducted tissue collection. ANJ, ANP and AA conducted data analysis.

## Data and materials availability

Upon acceptance of the paper, the authors plan to make all raw and processed data files publicly available via the Mendeley Data repository.

---

**Supplement Figure 1.**
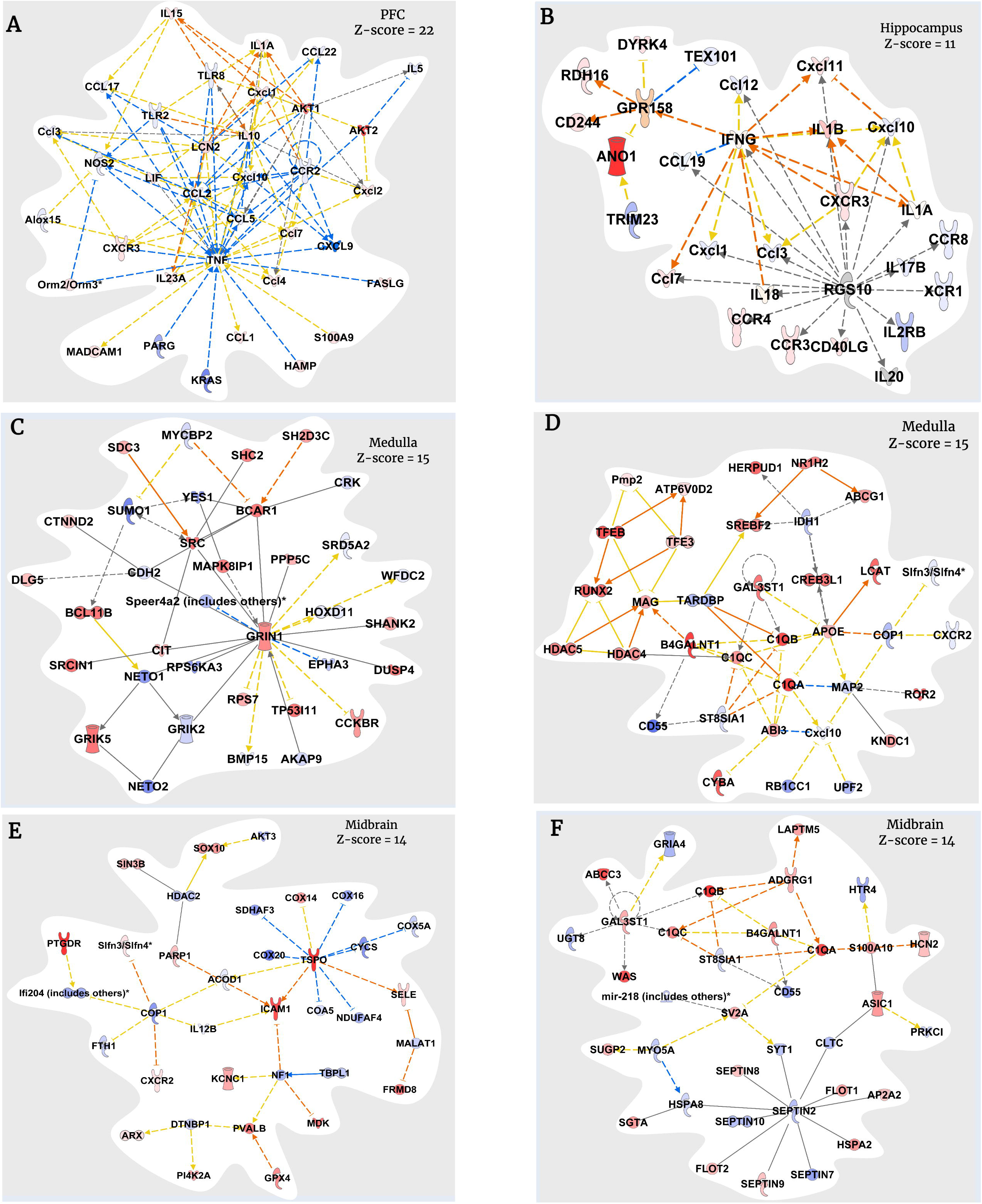
Top molecular networks (with significant Z scores) across different brain regions with genes linked to neuroinflammation and critical neuronal functions.

