## supplement for "Unraveling a comparative landscape of protein-coding genes linked to neuroimmune function during adulthood consequent of prenatal alcohol exposure": table1 anj030.docx

| **PFC**  **UPREGULATED** | | | **PFC**  **DOWNREGULATED** | | | **ACC**  **UPREGULATED** | | | | **ACC**  **DOWNREGULATED** | | | | |
| --- | --- | --- | --- | --- | --- | --- | --- | --- | --- | --- | --- | --- | --- | --- |
| **GENE** | **LOG2FC** | **P-ADJ** | **GENE** | **LOG2FC** | **P-ADJ** | **GENE** | **LOG2FC** | **P-ADJ** | | **GENE** | **LOG2FC** | | | **P-ADJ** |
| Pcsk1n | 0.34 | 6E-06 | Zfp965 | -0.48 | 2E-06 | Thsd4 | 0.37 | 8E-04 | | mt-Nd3 | -1.16 | | | 2E-25 |
| Chac1 | 0.33 | 3E-04 | Zfp942 | -0.41 | 3E-06 | Cela2a | 0.37 | 5E-03 | | mt-Atp8 | -1.00 | | | 6E-08 |
| Nkx3-1 | 0.32 | 2E-03 | Zfp943 | -0.39 | 3E-05 | Aldh4a1 | 0.27 | 6E-06 | | mt-Atp6 | -0.90 | | | 3E-19 |
| Ahnak2 | 0.32 | 4E-04 | Vbp1 | -0.37 | 1E-05 | AkA | 0.27 | 1E-03 | | mt-Nd4 | -0.73 | | | 9E-12 |
| Mmp11 | 0.32 | 4E-04 | mt-Nd3 | -0.36 | 1E-03 | Cgas | 0.26 | 1E-02 | | mt-Nd2 | -0.56 | | | 7E-08 |
| C1qtnf4 | 0.30 | 1E-03 | Ube3a | -0.34 | 3E-05 | Tsks | 0.25 | 1E-02 | | mt-Co2 | -0.46 | | | 2E-08 |
| C1qa | 0.30 | 4E-04 | Necab1 | -0.32 | 1E-03 |  |  |  | | mt-Co3 | -0.45 | | | 2E-07 |
| Smad6 | 0.29 | 3E-03 | Hnrnpu | -0.29 | 3E-04 |  |  |  | | mt-Cytb | -0.42 | | | 2E-06 |
| Fstl3 | 0.29 | 4E-03 | Cox20 | -0.28 | 3E-03 |  |  |  | | Nox1 | -0.38 | | | 1E-03 |
| Lama5 | 0.28 | 4E-04 | Syncrip | -0.27 | 4E-05 |  |  |  | | Ap1s2 | -0.37 | | | 5E-04 |
| Cd72 | 0.27 | 1E-03 | Klhl15 | -0.26 | 7E-04 |  |  |  | | Vma21 | -0.36 | | | 1E-05 |
| Prr7 | 0.26 | 4E-03 | Hnrnph1 | -0.26 | 2E-03 |  |  |  | | mt-Co1 | -0.35 | | | 1E-05 |
| Marveld1 | 0.26 | 2E-03 | Klhl28 | -0.26 | 2E-03 |  |  |  | | Ube3a | -0.32 | | | 7E-07 |
| H2-DMa | 0.26 | 4E-03 | Qki | -0.26 | 2E-03 |  |  | |  | mt-Nd4l | | -0.26 | 2E-03 | |
| Icam1 | 0.25 | 9E-03 | Stag2 | -0.25 | 2E-03 |  |  | |  | mt-Nd1 | | -0.25 | 5E-04 | |
| **HYPOTHALAMUS UPREGULATED** | | | **HYPOTHALAMUS DOWNREGULATED** | | | **HIPPOCAMPUS UPREGULATED** | | | | **HIPPOCAMPUS DOWNREGULATED** | | | | |
| **GENE** | **LOG2FC** | **P-ADJ** | **GENE** | **LOG2FC** | **P-ADJ** | **GENE** | **LOG2FC** | **P-ADJ** | | **GENE** | **LOG2FC** | | | **P-ADJ** |
| Mmp11 | 1.05 | 6E-12 | mt-Apt8 | -1.93 | 4E-09 | Crhr2 | 0.79 | 1E-03 | | mt-Atp8 | -1.10 | | | 6E-10 |
| N4bp3 | 1.01 | 5E-06 | Traf1 | -1.50 | 2E-04 | Thbs1 | 0.54 | 6E-03 | | mt-Nd3 | -1.09 | | | 2E-09 |
| Npas1 | 0.87 | 2E-04 | mt-Nd3 | -1.42 | 2E-09 | Mmp9 | 0.38 | 2E-02 | | mt-Atp6 | -0.85 | | | 3E-07 |
| Aldh3b1 | 0.74 | 3E-11 | mt-Apt6 | -1.34 | 4E-07 | Fzd7 | 0.38 | 9E-03 | | Gpr34 | -0.64 | | | 7E-08 |
| Irf2bp1 | 0.63 | 3E-06 | Cxcr4 | -1.26 | 3E-04 | Fzd8 | 0.37 | 1E-02 | | Samd9l | -0.63 | | | 2E-04 |
| Vamp1 | 0.60 | 4E-03 | mt-Nd4 | -1.19 | 4E-07 | Irf2bp1 | 0.36 | 2E-05 | | mt-Nd4 | -0.62 | | | 2E-04 |
| Slc4a2 | 0.59 | 7E-03 | Hba-a2 | -1.10 | 3E-03 | Il6ra | 0.31 | 1E-02 | | mt-Co2 | -0.52 | | | 2E-04 |
| Nat8l | 0.56 | 1E-03 | Hba-a1 | -1.09 | 3E-03 | H2-DMa | 0.30 | 8E-04 | | Htr2a | -0.43 | | | 1E-02 |
| Gja4 | 0.56 | 1E-03 | Unc5cl | -1.02 | 2E-03 | Mmp15 | 0.29 | 2E-02 | | Ncam2 | -0.41 | | | 4E-03 |
| Cox7c | 0.53 | 4E-04 | mt-Nd2 | -0.91 | 5E-05 | Cxcr4 | 0.29 | 4E-02 | | Grm3 | -0.34 | | | 2E-02 |
| Pvalb | 0.52 | 1E-02 | Neurod6 | -0.86 | 3E-03 | Mmp14 | 0.28 | 1E-02 | | Jak2 | -0.33 | | | 1E-02 |
| Kcnj11 | 0.52 | 4E-03 | Nlrp12 | -0.78 | 4E-03 | Atf6b | 0.27 | 2E-02 | | Gria4 | -0.32 | | | 2E-02 |
| H1f3 | 0.49 | 7E-03 | mt-Cytb | -0.73 | 1E-04 | Gas6 | 0.27 | 2E-02 | | Nlrp6 | -0.29 | | | 3E-02 |
| Olig1 | 0.48 | 4E-03 | Bace2 | -0.69 | 2E-03 | Gas1 | 0.27 | | 4E-02 | Bcl2 | | -0.27 | 5E-02 | |
| Cd72 | 0.45 | 1E-03 | Aqp4 | -0.53 | 3E-04 | Hspa1b | 0.26 | | 4E-02 | Nr3c1 | | -0.25 | 8E-03 | |
| **MEDULLA**  **UPREGULATED** | | | **MEDULLA DOWNREGULATED** | | | **MIDBRAIN**  **UPREGULATED** | | | | **MIDBRAIN DOWNREGULATED** | | | | |
| **GENE** | **LOG2FC** | **P-ADJ** | **GENE** | **LOG2FC** | **P-ADJ** | **GENE** | **LOG2FC** | **P-ADJ** | | **GENE** | **LOG2FC** | | | **P-ADJ** |
| Lyve1 | 1.26 | 6E-06 | mt-Atp8 | -1.64 | 5E-22 | Wnt10a | 0.81 | 1E-04 | | mt-Atp8 | -2.07 | | | 2E-21 |
| H3c3 | 0.77 | 3E-04 | mt-Atp6 | -1.34 | 5E-24 | Ier2 | 0.72 | 5E-09 | | mt-Nd3 | -1.67 | | | 3E-16 |
| H3c2 | 0.76 | 2E-03 | mt-Nd3 | -1.26 | 8E-36 | Gas1 | 0.58 | 1E-08 | | mt-Atp6 | -1.59 | | | 1E-12 |
| Cebpb | 0.63 | 4E-04 | mt-Nd4 | -1.11 | 8E-35 | Jund | 0.56 | 2E-10 | | mt-Nd4 | -1.44 | | | 2E-11 |
| H3c4 | 0.59 | 4E-04 | mt-Co2 | -0.96 | 1E-29 | Ifi27l2b | 0.55 | 2E-03 | | mt-Nd2 | -1.32 | | | 4E-12 |
| Nfatc4 | 0.59 | 1E-03 | mt-Cytb | -0.96 | 6E-22 | Irf2bp1 | 0.55 | 4E-11 | | mt-Cytb | -1.07 | | | 1E-08 |
| Jund | 0.52 | 4E-04 | Gpr34 | -0.91 | 2E-17 | Ccl21d | 0.51 | 7E-04 | | mt-Nd1 | -1.05 | | | 7E-11 |
| Nfkbil1 | 0.52 | 3E-03 | mt-Co3 | -0.90 | 3E-19 | Itga11 | 0.51 | 5E-04 | | mt-Co2 | -1.01 | | | 2E-10 |
| Irfbp1 | 0.51 | 4E-03 | mt-Nd2 | -0.90 | 4E-22 | Il17rc | 0.50 | 2E-04 | | mt-Co3 | -1.00 | | | 2E-22 |
| Pcsk1n | 0.51 | 4E-04 | Nox1 | -0.83 | 4E-12 | Ccl21a | 0.44 | 2E-03 | | Nox1 | -0.74 | | | 6E-07 |
| C1qtnf4 | 0.51 | 1E-03 | mt-Nd1 | -0.76 | 1E-17 | Ccl21b | 0.44 | 3E-03 | | Gvin1 | -0.64 | | | 3E-11 |
| Il34 | 0.31 | 4E-03 | Cd36 | -0.75 | 2E-04 | Tnfrsf4 | 0.33 | 2E-02 | | Atp11c | -0.62 | | | 2E-10 |
| C1qtnf12 | 0.31 | 2E-03 | Ube3a | -0.65 | 4E-36 | Il1r1 | 0.33 | 2E-02 | | Il2 | -0.61 | | | 1E-04 |
| Il11ra2 | 0.31 | 2E-02 | Vma21 | -0.62 | 7E-23 | Il16 | 0.30 | 2E-02 | | Il1rapl1 | -0.55 | | | 2E-08 |
| Il21r | 0.29 | 2E-02 | Cox20 | -0.51 | 3E-06 | Ccl19 | 0.29 | 3E-02 | | Il4 | -0.27 | | | 2E-02 |

**Table 1. Top significant DEGs (with a minimum of log2FC of 0.25, selected from the top 15) linked to neuroimmune function.**

| **PFC**  **UPREGULATED** | | | **PFC**  **DOWNREGULATED** | | | **ACC**  **UPREGULATED** | | | | **ACC**  **DOWNREGULATED** | | | | |
| --- | --- | --- | --- | --- | --- | --- | --- | --- | --- | --- | --- | --- | --- | --- |
| **GENE** | **LOG2FC** | **P-ADJ** | **GENE** | **LOG2FC** | **P-ADJ** | **GENE** | **LOG2FC** | **P-ADJ** | | **GENE** | **LOG2FC** | | | **P-ADJ** |
| Pcsk1n | 0.34 | 6E-06 | Zfp965 | -0.48 | 2E-06 | Thsd4 | 0.37 | 8E-04 | | mt-Nd3 | -1.16 | | | 2E-25 |
| Chac1 | 0.33 | 3E-04 | Zfp942 | -0.41 | 3E-06 | Cela2a | 0.37 | 5E-03 | | mt-Atp8 | -1.00 | | | 6E-08 |
| Nkx3-1 | 0.32 | 2E-03 | Zfp943 | -0.39 | 3E-05 | Aldh4a1 | 0.27 | 6E-06 | | mt-Atp6 | -0.90 | | | 3E-19 |
| Ahnak2 | 0.32 | 4E-04 | Vbp1 | -0.37 | 1E-05 | AkA | 0.27 | 1E-03 | | mt-Nd4 | -0.73 | | | 9E-12 |
| Mmp11 | 0.32 | 4E-04 | mt-Nd3 | -0.36 | 1E-03 | Cgas | 0.26 | 1E-02 | | mt-Nd2 | -0.56 | | | 7E-08 |
| C1qtnf4 | 0.30 | 1E-03 | Ube3a | -0.34 | 3E-05 | Tsks | 0.25 | 1E-02 | | mt-Co2 | -0.46 | | | 2E-08 |
| C1qa | 0.30 | 4E-04 | Necab1 | -0.32 | 1E-03 |  |  |  | | mt-Co3 | -0.45 | | | 2E-07 |
| Smad6 | 0.29 | 3E-03 | Hnrnpu | -0.29 | 3E-04 |  |  |  | | mt-Cytb | -0.42 | | | 2E-06 |
| Fstl3 | 0.29 | 4E-03 | Cox20 | -0.28 | 3E-03 |  |  |  | | Nox1 | -0.38 | | | 1E-03 |
| Lama5 | 0.28 | 4E-04 | Syncrip | -0.27 | 4E-05 |  |  |  | | Ap1s2 | -0.37 | | | 5E-04 |
| Cd72 | 0.27 | 1E-03 | Klhl15 | -0.26 | 7E-04 |  |  |  | | Vma21 | -0.36 | | | 1E-05 |
| Prr7 | 0.26 | 4E-03 | Hnrnph1 | -0.26 | 2E-03 |  |  |  | | mt-Co1 | -0.35 | | | 1E-05 |
| Marveld1 | 0.26 | 2E-03 | Klhl28 | -0.26 | 2E-03 |  |  |  | | Ube3a | -0.32 | | | 7E-07 |
| H2-DMa | 0.26 | 4E-03 | Qki | -0.26 | 2E-03 |  |  | |  | mt-Nd4l | | -0.26 | 2E-03 | |
| Icam1 | 0.25 | 9E-03 | Stag2 | -0.25 | 2E-03 |  |  | |  | mt-Nd1 | | -0.25 | 5E-04 | |
| **HYPOTHALAMUS UPREGULATED** | | | **HYPOTHALAMUS DOWNREGULATED** | | | **HIPPOCAMPUS UPREGULATED** | | | | **HIPPOCAMPUS DOWNREGULATED** | | | | |
| **GENE** | **LOG2FC** | **P-ADJ** | **GENE** | **LOG2FC** | **P-ADJ** | **GENE** | **LOG2FC** | **P-ADJ** | | **GENE** | **LOG2FC** | | | **P-ADJ** |
| Mmp11 | 1.05 | 6E-12 | mt-Apt8 | -1.93 | 4E-09 | Crhr2 | 0.79 | 1E-03 | | mt-Atp8 | -1.10 | | | 6E-10 |
| N4bp3 | 1.01 | 5E-06 | Traf1 | -1.50 | 2E-04 | Thbs1 | 0.54 | 6E-03 | | mt-Nd3 | -1.09 | | | 2E-09 |
| Npas1 | 0.87 | 2E-04 | mt-Nd3 | -1.42 | 2E-09 | Mmp9 | 0.38 | 2E-02 | | mt-Atp6 | -0.85 | | | 3E-07 |
| Aldh3b1 | 0.74 | 3E-11 | mt-Apt6 | -1.34 | 4E-07 | Fzd7 | 0.38 | 9E-03 | | Gpr34 | -0.64 | | | 7E-08 |
| Irf2bp1 | 0.63 | 3E-06 | Cxcr4 | -1.26 | 3E-04 | Fzd8 | 0.37 | 1E-02 | | Samd9l | -0.63 | | | 2E-04 |
| Vamp1 | 0.60 | 4E-03 | mt-Nd4 | -1.19 | 4E-07 | Irf2bp1 | 0.36 | 2E-05 | | mt-Nd4 | -0.62 | | | 2E-04 |
| Slc4a2 | 0.59 | 7E-03 | Hba-a2 | -1.10 | 3E-03 | Il6ra | 0.31 | 1E-02 | | mt-Co2 | -0.52 | | | 2E-04 |
| Nat8l | 0.56 | 1E-03 | Hba-a1 | -1.09 | 3E-03 | H2-DMa | 0.30 | 8E-04 | | Htr2a | -0.43 | | | 1E-02 |
| Gja4 | 0.56 | 1E-03 | Unc5cl | -1.02 | 2E-03 | Mmp15 | 0.29 | 2E-02 | | Ncam2 | -0.41 | | | 4E-03 |
| Cox7c | 0.53 | 4E-04 | mt-Nd2 | -0.91 | 5E-05 | Cxcr4 | 0.29 | 4E-02 | | Grm3 | -0.34 | | | 2E-02 |
| Pvalb | 0.52 | 1E-02 | Neurod6 | -0.86 | 3E-03 | Mmp14 | 0.28 | 1E-02 | | Jak2 | -0.33 | | | 1E-02 |
| Kcnj11 | 0.52 | 4E-03 | Nlrp12 | -0.78 | 4E-03 | Atf6b | 0.27 | 2E-02 | | Gria4 | -0.32 | | | 2E-02 |
| H1f3 | 0.49 | 7E-03 | mt-Cytb | -0.73 | 1E-04 | Gas6 | 0.27 | 2E-02 | | Nlrp6 | -0.29 | | | 3E-02 |
| Olig1 | 0.48 | 4E-03 | Bace2 | -0.69 | 2E-03 | Gas1 | 0.27 | | 4E-02 | Bcl2 | | -0.27 | 5E-02 | |
| Cd72 | 0.45 | 1E-03 | Aqp4 | -0.53 | 3E-04 | Hspa1b | 0.26 | | 4E-02 | Nr3c1 | | -0.25 | 8E-03 | |
| **MEDULLA**  **UPREGULATED** | | | **MEDULLA DOWNREGULATED** | | | **MIDBRAIN**  **UPREGULATED** | | | | **MIDBRAIN DOWNREGULATED** | | | | |
| **GENE** | **LOG2FC** | **P-ADJ** | **GENE** | **LOG2FC** | **P-ADJ** | **GENE** | **LOG2FC** | **P-ADJ** | | **GENE** | **LOG2FC** | | | **P-ADJ** |
| Lyve1 | 1.26 | 6E-06 | mt-Atp8 | -1.64 | 5E-22 | Wnt10a | 0.81 | 1E-04 | | mt-Atp8 | -2.07 | | | 2E-21 |
| H3c3 | 0.77 | 3E-04 | mt-Atp6 | -1.34 | 5E-24 | Ier2 | 0.72 | 5E-09 | | mt-Nd3 | -1.67 | | | 3E-16 |
| H3c2 | 0.76 | 2E-03 | mt-Nd3 | -1.26 | 8E-36 | Gas1 | 0.58 | 1E-08 | | mt-Atp6 | -1.59 | | | 1E-12 |
| Cebpb | 0.63 | 4E-04 | mt-Nd4 | -1.11 | 8E-35 | Jund | 0.56 | 2E-10 | | mt-Nd4 | -1.44 | | | 2E-11 |
| H3c4 | 0.59 | 4E-04 | mt-Co2 | -0.96 | 1E-29 | Ifi27l2b | 0.55 | 2E-03 | | mt-Nd2 | -1.32 | | | 4E-12 |
| Nfatc4 | 0.59 | 1E-03 | mt-Cytb | -0.96 | 6E-22 | Irf2bp1 | 0.55 | 4E-11 | | mt-Cytb | -1.07 | | | 1E-08 |
| Jund | 0.52 | 4E-04 | Gpr34 | -0.91 | 2E-17 | Ccl21d | 0.51 | 7E-04 | | mt-Nd1 | -1.05 | | | 7E-11 |
| Nfkbil1 | 0.52 | 3E-03 | mt-Co3 | -0.90 | 3E-19 | Itga11 | 0.51 | 5E-04 | | mt-Co2 | -1.01 | | | 2E-10 |
| Irfbp1 | 0.51 | 4E-03 | mt-Nd2 | -0.90 | 4E-22 | Il17rc | 0.50 | 2E-04 | | mt-Co3 | -1.00 | | | 2E-22 |
| Pcsk1n | 0.51 | 4E-04 | Nox1 | -0.83 | 4E-12 | Ccl21a | 0.44 | 2E-03 | | Nox1 | -0.74 | | | 6E-07 |
| C1qtnf4 | 0.51 | 1E-03 | mt-Nd1 | -0.76 | 1E-17 | Ccl21b | 0.44 | 3E-03 | | Gvin1 | -0.64 | | | 3E-11 |
| Il34 | 0.31 | 4E-03 | Cd36 | -0.75 | 2E-04 | Tnfrsf4 | 0.33 | 2E-02 | | Atp11c | -0.62 | | | 2E-10 |
| C1qtnf12 | 0.31 | 2E-03 | Ube3a | -0.65 | 4E-36 | Il1r1 | 0.33 | 2E-02 | | Il2 | -0.61 | | | 1E-04 |
| Il11ra2 | 0.31 | 2E-02 | Vma21 | -0.62 | 7E-23 | Il16 | 0.30 | 2E-02 | | Il1rapl1 | -0.55 | | | 2E-08 |
| Il21r | 0.29 | 2E-02 | Cox20 | -0.51 | 3E-06 | Ccl19 | 0.29 | 3E-02 | | Il4 | -0.27 | | | 2E-02 |
