## supplement for "Unraveling a comparative landscape of protein-coding genes linked to neuroimmune function during adulthood consequent of prenatal alcohol exposure": table 2 anj030.docx

| **Table 2: Selected overlapping significant DEGs across brain regions.** | | | | | | | | | | | |
| --- | --- | --- | --- | --- | --- | --- | --- | --- | --- | --- | --- |
| **Genes** | **MEDULLA** | | **MIDBRAIN** | | **HYPOTHALAMUS** | | **HIPPOCAMPUS** | | **ACC** | | **PFC** |
| **Medulla_Midbrain** |  | |  | |  | |  | |  | |  |
| Irf2bpl | 0.36 | | 0.49 | | - | | - | | - | | - |
| Dnajb9 | -0.26 | | -0.35 | | - | | - | | - | | - |
| Akt3 | -0.28 | | -0.29 | | - | | - | | - | | - |
| Cd164 | -0.46 | | -0.47 | | - | | - | | - | | - |
| Itga4 | -0.27 | | -0.29 | | - | | - | | - | | - |
| Picalm | -0.3 | | -0.32 | | - | | - | | - | | - |
| Ccl21a | 0.36 | | 0.44 | | - | | - | | - | | - |
| Hspd1 | -0.25 | | -0.3 | | - | | - | | - | | - |
| Il2 | -0.38 | | -0.61 | | - | | - | | - | | - |
| Wnt10a | 0.44 | | 0.81 | | - | | - | | - | | - |
| Hspa4l | -0.25 | | -0.28 | | - | | - | | - | | - |
| C1qtnf1 | 0.29 | | 0.37 | | - | | - | | - | | - |
| Tlr3 | -0.4 | | -0.46 | | - | | - | | - | | - |
| Atp11c | -0.64 | | -0.62 | | - | | - | | - | | - |
| Il34 | 0.31 | | 0.32 | | - | | - | | - | | - |
| **Hypothalamus_Hippocampus** |  | |  | |  | |  | |  | |  |
| Gas6 | - | | - | | 0.36 | | 0.27 | | - | | - |
| Nr3c1 | - | | - | | -0.32 | | -0.25 | | - | | - |
| Npas1 | - | | - | | 0.87 | | 0.26 | | - | | - |
| Cxcr4 | - | | - | | -1.26 | | 0.29 | | - | | - |
| **Medulla_Midbrain_Hypothalamus** |  | |  | |  | |  | |  | |  |
| Ier3 | -0.44 | | 0.27 | | 0.39 | | - | | - | | - |
| Ift88 | -0.25 | | -0.25 | | -0.3 | | - | | - | | - |
| Pten | -0.27 | | -0.32 | | -0.26 | | - | | - | | - |
| Sirt1 | -0.45 | | -0.34 | | -0.49 | | - | | - | | - |
| Dnaja1 | -0.36 | | -0.38 | | -0.36 | | - | | - | | - |
| C1qb | 0.35 | | 0.43 | | 0.33 | | - | | - | | - |
| Irf3 | 0.31 | | 0.27 | | 0.34 | | - | | - | | - |
| Gja4 | 0.29 | | 0.33 | | 0.56 | | - | | - | | - |
| Hsp90aa1 | -0.32 | | -0.29 | | -0.28 | | - | | - | | - |
| **Medulla_Midbrain_PFC** |  | |  | |  | |  | |  | |  |
| C1qtnf4 | 0.51 | | 0.48 | | - | | - | | - | | 0.30 |
| Prr7 | 0.36 | | 0.34 | | - | | - | | - | | 0.26 |
| C1qa | 0.36 | | 0.39 | | - | | - | | - | | 0.30 |
| Icam1 | 0.28 | | 0.38 | | - | | - | | - | | 0.25 |
| **Medulla_Hypothalamus_Hippocampus** |  | |  | |  | |  | |  | |  |
| Dnajb4 | -0.33 | | - | | -0.28 | | -0.25 | | - | | - |
| **Midbrain_Hypothalamus_Hippocampus** |  | |  | |  | |  | |  | |  |
| Mmp15 | - | | 0.27 | | 0.52 | | 0.29 | | - | | - |
| **Medulla_Midbrain_Hypothalamus_Hippocampus** |  |  | |  | |  | |  | |  | |
| mt-Nd5 | -0.37 | -0.68 | | -0.44 | | -0.26 | | - | | - | |
| Cdc73 | -0.48 | -0.29 | | -0.5 | | -0.29 | | - | | - | |
| Itgbl1 | -0.48 | -0.44 | | -0.57 | | -0.37 | | - | | - | |
| Irf2bp1 | 0.51 | 0.55 | | 0.63 | | 0.36 | | - | | - | |
| Gpr34 | -0.91 | -0.72 | | -0.69 | | -0.64 | | - | | - | |
| Jak2 | -0.33 | -0.26 | | -0.3 | | -0.33 | | - | | - | |
| **Medulla_Midbrain_Hypothalamus_PFC** |  |  | |  | |  | |  | |  | |
| Hnrnph1 | -0.31 | -0.29 | | -0.29 | | - | | - | | -0.26 | |
| Cd72 | 0.41 | 0.34 | | 0.45 | | - | | - | | 0.27 | |
| Mmp11 | 0.5 | 0.52 | | 1.05 | | - | | - | | 0.32 | |
| Cox20 | -0.51 | -0.48 | | -0.55 | | - | | - | | -0.28 | |
| Jund | 0.52 | 0.56 | | 0.45 | | - | | - | | 0.28 | |
| **Medulla_Midbrain_Hippocampus_PFC** |  |  | |  | |  | |  | |  | |
| Marveld1 | 0.26 | 0.27 | | - | | -0.33 | | - | | 0.26 | |
| Qki | -0.44 | -0.44 | | - | | -0.32 | | - | | -0.26 | |
| **Medulla_Hypothalamus_Hippocampus_PFC** |  |  | |  | |  | |  | |  | |
| Chac1 | 0.37 | - | | 0.34 | | 0.31 | | - | | 0.33 | |
| Arglu1 | -0.25 | - | | -0.28 | | -0.27 | | - | | -0.25 | |
| **Medulla_Midbrain_Hypothalamus_Hippocampus_PFC** |  |  | |  | |  | |  | |  | |
| Hnrnpu | -0.45 | -0.43 | | -0.47 | | -0.26 | | - | | -0.29 | |
| **Medulla_Midbrain_Hypothalamus_Hippocampus_ACC** |  |  | |  | |  | |  | |  | |
| Nox1 | -0.83 | -0.74 | | -0.7 | | -0.43 | | -0.38 | | - | |
| mt-Co2 | -0.96 | -1.01 | | -0.67 | | -0.52 | | -0.46 | | - | |
| mt-Co3 | -0.9 | -1.00 | | -0.60 | | -0.44 | | -0.45 | | - | |
| mt-Nd2 | -0.9 | -1.32 | | -0.91 | | -0.48 | | -0.56 | | - | |
| mt-Nd4l | -0.72 | -0.99 | | -0.79 | | -0.25 | | -0.26 | | - | |
| mt-Nd1 | -0.76 | -1.05 | | -0.61 | | -0.42 | | -0.25 | | - | |
| mt-Co1 | -0.54 | -0.68 | | -0.38 | | -0.32 | | -0.35 | | - | |
| mt-Cytb | -0.96 | -1.07 | | -0.73 | | -0.41 | | -0.42 | | - | |
| mt-Nd4l | -0.72 | -0.99 | | -0.79 | | -0.25 | | -0.26 | | - | |
| mt-Atp8 | -1.64 | -2.07 | | -1.93 | | -1.10 | | -1.00 | | - | |
| **Medulla_Midbrain_Hypothalamus_Hippocampus_ACC_PFC** |  |  | |  | |  | |  | |  | |
| Ube3a | -0.65 | -0.72 | | -0.52 | | -0.39 | | -0.32 | | -0.34 | |
| Vma21 | -0.62 | -0.60 | | -0.47 | | -0.31 | | -0.36 | | -0.30 | |
| mt-Nd3 | -1.26 | -1.67 | | -1.42 | | -1.09 | | -1.16 | | -0.36 | |

*Log_2_FCs are shown for each significant DEG in the indicated brain region. Blank cells indicate that the gene did not meet dysregulation criteria in that region (log_2_FC > |0.25| *p-adj* < 0.05).
